# H^+^ drives ultra-fast root-to-root responses to wounding

**DOI:** 10.1101/2024.10.04.616436

**Authors:** Julie Ducla, Luciano Di Fino, Andriani Mentzelopoulou, Loïc Talide, Maarten Besten, Iwona Bernacka-Wojcik, Theresia Arbring Sjöström, Nageena Zahid, Sandra Jämtgård, Eleni Stavrinidou, Totte Niittylä, Joris Sprakel, Peter Marhavý

## Abstract

The plant-to-plant communication of damage is vital for plants to mount pre-emptive defensive responses in the face of threats. A variety of threats to the well-being of plants are found below ground; yet how plant roots activate inter-plant communication is largely unclear. Here we demonstrate that a wounded root rapidly releases protons (H^+^), that travel faster than any other “known” soluble biochemical signal due to a specialised diffusion mechanism. Within seconds after damage, cells in neighbouring unwounded roots sense the acidification and activate tissue-specific Ca^2+^ damage signalling. In turn, this triggers a differential growth response allowing the unwounded root to avoid the site of a potential threat. Our results reveal a non-canonical rapid response mechanism for inter-plant communication based on ultrafast proton diffusion.

## Main/Introduction

While animals have evolved complex communication systems to enhance their fitness in response to threats**^1,2,3^**, plants, lacking such overt communicative abilities, also exhibit fascinating forms of inter-plant communication**^4,5,6^**. The evolutionary benefit for neighboring plants to detect the signal of a close-by attack is obvious, as it can enable plants to mobilize their resources and get prepared for a potential attack and increase population level fitness/survival. Hence, it is clear that molecular events linked to an early response are critical for the survival of plants. Although several molecular regulators have been identified in defensive stress response signaling, the exact mechanisms by which such signals are perceived are still largely unknown**^7,8,9,10^**. In aerial tissues, communication systems for long distance plant-to-plant signaling have been extensively investigated. Biochemical and genetic evidence have revealed changes induced in plants responding to volatiles released by other plants attacked by pathogens or herbivores**^11,5,12,13^**. It has been proposed that below ground, plants share chemical informations**^14,15,16,17^** which can help them to communicate about various conditions**^18,19^**. These signals can prompt neighboring plants to enhance their physical defenses, such as strengthening cell walls, or to increase the production of secondary metabolites that deter herbivores and inhibit pathogen growth**^20^**. It has been shown that the level of specific root exudates plays a crucial role in both self-recognition and recognition of neighboring plants**^21^**. Specifically, the accumulation of compounds secreted by the same species inhibits root growth**^22,23,24^**. Responses to nutrients can also dominate over the avoidance of neighboring plants, and that plants integrate signals from both nutrients and competitors to fine-tune their growth and development**^25^**. When looking at proton (H^+^) fluxes in the root, which are important for root growth, it was found that the response to environmental stresses like adverse soil conditions such as drought, salinity, hypoxia, and nutrient deficiencies relies heavily on the regulation of H^+^ fluxes at the root tip**^26^**. Furthermore, protons may act as signaling intermediates, particularly during apoplastic pH shifts caused by immunological responses**^27^**. The growth-immunity balance is maintained by the equilibrium of apoplastic pH, with the changes associated with pattern-triggered immunity (PTI) referred to as the “root alkaline switch”**^27^**.

Here, we report that a wounded root releases a very rapid signal to neighbouring roots and activates Ca^2+^ signalling in the unwounded roots within seconds. Through modelling and kinetics response analyses, we conclude that all conventional soluble signals (ions, hormones, peptides) diffuse too slowly to explain the ultrafast root-to-root communication. We show instead that protons (H^+^) are the causative signal: the wounded cell releases a wave of protons that travels through the medium an order-of-magnitude faster than any other soluble signal by the so-called proton hopping mechanism. The resulting acidification of the medium and the cell walls of neighbouring undamaged roots is quickly detected activating a cortex-specific Ca^2+^ signal. In turn, the unwounded root displays differential cell elongation in the root apical meristem, allowing the undamaged root to bend away from the source of potential threat as a damage avoidance mechanism. We reveal a previously unacknowledged mechanism by which protons act as signals in inter-plant communication that can operate on the scale of seconds, much faster than conventional signalling pathways. Such rapid responses can explain how plants quickly adapt to environmental stressors and provide a significant advantage in fitness and survival, allowing them to better defend against threats and optimize resource allocation.

## Results

### Root-to-root wounding perception was visualised using a genetically encoded calcium sensor

A well-established early response to damage is the emission of a wave of cytosolic calcium, [Ca^2+^]_cyt_, that spreads throughout the plant tissue from the site of damage (Extended Data Fig. 1a). This can be probed using calcium sensing reporter lines, regardless of whether localised or systemic signaling is being studied^28^. We used the intensiometric calcium reporter lines, R-GECO1^29^ and GCaMP3^30^, to visualise wound sensing from one root, the emitter (E), which is damaged, to a neighbour, that is undamaged, and acts as a receiver (R). This was accomplished by single-cell laser ablation^31,32^ of an epidermal cell in the E, followed by monitoring the Ca^2+^ response in the unwounded R (Fig. 1a-d; Movie S2). Upon wounding the E root, both the wounded^31^ E and the R root rapidly activated a Ca^2+^ response (Extended Data Fig. 1a,d) characteristic of a wounding stress, even though the R root was not wounded (Fig. 1c, Movie S2). Clearly, there is both internal and inter-plant damage perception (Movie S1). Interestingly, the emitted signal is not *Arabidopsis* specific; ablating a single epidermal cell in *Solanum lycopersicum* E roots, led to the same response in a R-GECO1 expressing *Arabidopsis thaliana* receiver root (Fig. 1f-g; Movie S2) as in the experiment between two *Arabidopsis* roots (Fig. 1c-d; Extended Data Fig. 1a-d). This suggests that the sensed signal is generic.

**Figure 1:**
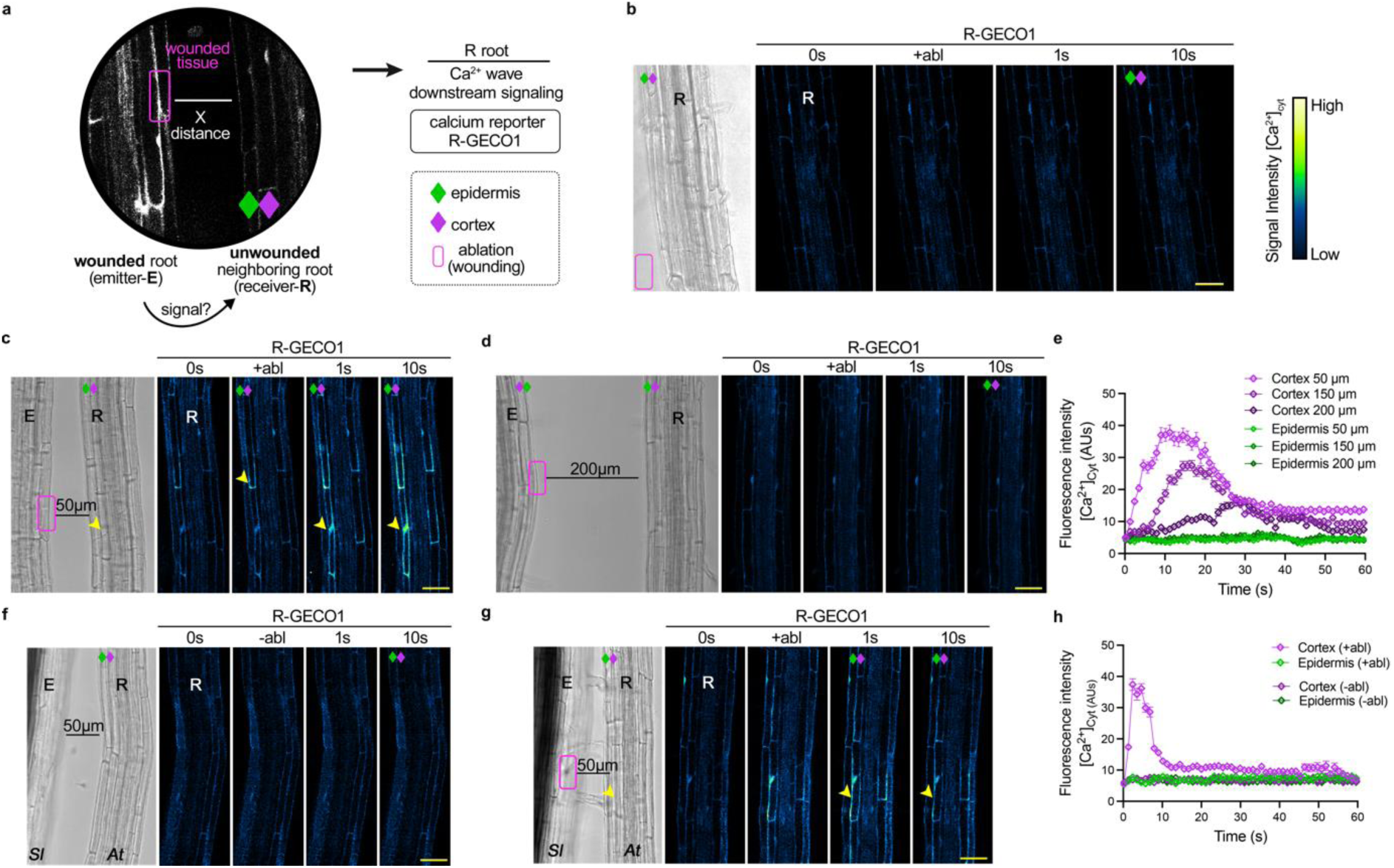
Harnessing the Ca^2+^ wave to visualise and understand plant wounding. a) Schematic overview of the wound induced root communication concept described in this paper. Emitter (E) root (i.e. wounded root; magenta boxes indicate the ablated/wounded area) emits a signal, which is perceived from the Receiver (R) root. *At*R-GECO1 tensiometric calcium reporter line (Green-Fire-Blue) is used as a downstream signalling read-out. Epidermis and cortex cell types are indicated by green and purple diamonds, respectively. b) Representative time-lapse imaging of *At*R-GECO1 over a 60-second time interval following laser ablation in the medium close to the root. Ablation of the solid agar medium did not induce any Ca^2+^ response. c) Single-cell laser ablation of the epidermal cell of the E root expressing *At*R-GECO1 triggers an immediate, cortex-specific Ca^2+^ wave when at a distance of 50 μm from the neighbouring R root. d) No Ca^2+^ wave was observed when the E root was 200 μm from the R root. e) Quantification of cortex-specific fluorescence intensity of cytosolic calcium ([Ca^2+^]_cyt_ in AUs) from (b-d) when the E root was ablated at distances between 50 μm-200 μm from the R root (n=10, data represented as mean ± SD). F-g) Single-cell laser ablation of a *Solanum lycopersicum* root E induced an immediate cortical Ca^2+^ response in the neighbouring *At*R-GECO1 expressing roots R. h) Quantification of e) (n=10, data represented as mean ± SD). The images in b, c, d, f and g all represent results from three independent biological replicates (N=3). The images (Green Fire Blue) only display the signal perceived in the receiver R root to facilitate the observation (complete video of (c) and (g) are available as Videos S1 and S2, respectively). The scale bars for a-e) are 50 μm. Green diamonds indicate the epidermal cell layer, while purple diamonds indicate the cortex. Magenta boxes indicate the ablation site. Yellow arrows indicate the calcium wave response. The signal intensity of the R-GECO1 calcium reporter (Green-Fire-Blue) is visualized according to the spectrum in 1a. AUs; Arbitrary Units.

To confirm that this is a wounding response and not an artefact due to the laser ablation, e.g. due to photothermal heating, we performed a control experiment in which we ablated/heated the medium in close proximity to a receiver root (Fig. 1a, Extended data1, Movie S3). Here, we observed no increase in cytosolic calcium concentrations ([Ca^2+^]_cyt_).

Based on these observations, we hypothesize that some fast-diffusing, soluble biochemical signal spreads from the E to the R root through the aqueous medium. We thus tested if ablation of an epidermal cell in the E root, facing away from the root R would induce a calcium wave in the R (Extended Data Fig. 1b). Here, we observed no change in cytosolic calcium in the R roots. This hints at a rapid “dilution” of the signal in the area between the face of the root that is damaged and the surroundings. This suggests that the signal is not effectively transmitted laterally through the tissue to another side. Instead, it appears that the proximity of the damaged cells is crucial for initiating the response, with the signal being rapidly diluted as it moves away from the site of damage. We therefore systematically tested the effect of root-to-root distance in the capacity of receiver roots to pick up the warning signal (Fig. 1e). We found that the capacity of R roots to sense the damage gradually decreased with increasing distance and appears to be lost when roots are further apart than 200 μm. This supports the notion of a diffusible compound working on short distance with a limit of 200 μm as the causative warning signal.

Next, we asked what part of the R root is most susceptible to perceiving the damage signal. We ablated epidermal cells of the E root facing various zones of the R root (Fig. 2). We find that the strongest increase in cytoplasmic calcium occurred in the cortical cell file of the differentiated zone (DZ) of the root, rather than the cortical cell file associated with the elongation zone (EZ) or apical meristematic zone (AMZ) of the root, areas which demonstrated a more even distribution of ([Ca^2+^]_cyt_) across all cell files (Fig. 2). This suggests that plant roots have the ability to detect and respond to even a single-cell wound in neighbouring plants in a tissue-specific manner (Movie S1).

**Figure 2:**
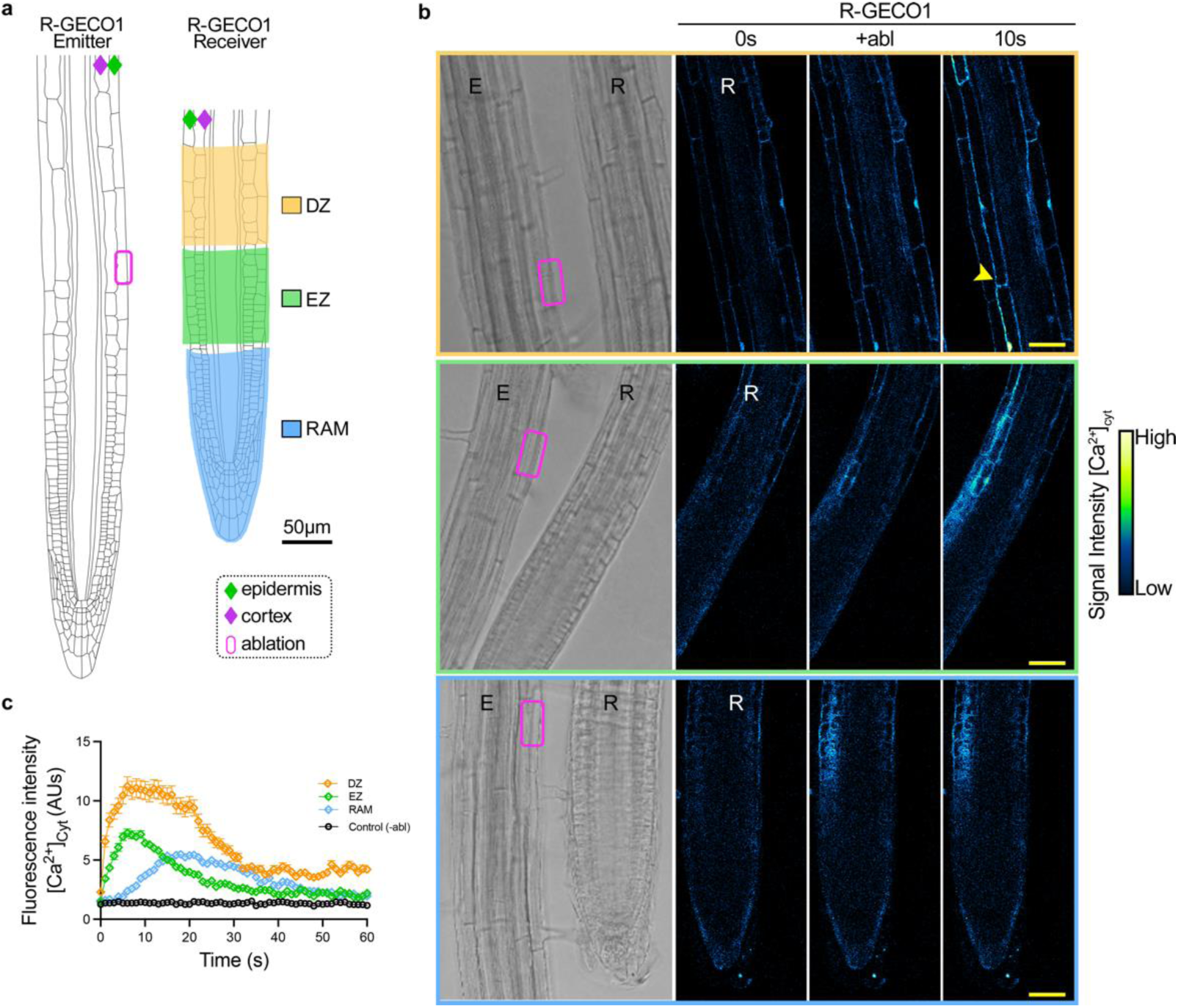
A Ca^2+^ wave at the cortex is initiated in the differentiated zone of the root in response to the wounding of neighbouring root. a) Schematic illustration of the different root zones used for [Ca^2+^]_cyt_ quantification and visualization in the receiver R root. c) Visualisation of the cortex-specific Ca^2+^ wave (using Green-Fire-Blue) over time in the R root, which includes the differentiated zone (DZ), elongation zone (EZ), and root apical meristematic zone (RAM). In the MZ, the cortex-specific Ca^2+^ wave initiates in the differentiated zone, while the Ca^2+^ wave is present in all cell files in the EZ and AMZ. d) Graph showing the cytosolic Ca^2+^ concentrations ([Ca^2+^]_cyt_) over time, presented as fluorescence intensity (AUs), from panel (c) in the R root. Data are represented as mean ± SD, with n=10. The graph in c represents the results from three independent biological replicates (N=3). a-c) scale bars, 50 μm; green diamonds indicate the epidermal cell layer, while purple diamonds indicate the cortex. Magenta boxes indicate the ablation site and the yellow arrows the calcium wave. Yellow arrows indicate the calcium wave response. The signal intensity of the R-GECO1 calcium reporter (Green-Fire-Blue) is visualized according to the spectrum in 2b. AUs; Arbitrary Units.

### Rapid H^+^ release during wounding is responsible for the root-to-root communication

Previous research has demonstrated that *in planta* electrical and hydraulic activity^33,34^ can be used to measure the speed of wound sensing. The induction of the [Ca2^+^]_cyt_ signal in R roots after wounding E happens within one second (Fig. 1e, h and Fig. 2c). According to this time-frame, along with the Grotthus mechanism of proton diffusion, we hypothesise that mobile H^+^ protons can elicit a significantly faster response than even the smallest non-hydrogen ions^35^, and consequently, could be responsible for root-to-root signaling (Extended Data Table 1). To test this hypothesis, we first calculated the diffusive time scale (τ_D_) for the different classes of compounds that are potentially released during wounding according to the surface distances (Extended Data Table 1). The calculations revealed that protons can in theory reach the neighbouring root much faster than other compounds, requiring between 0.3 s to 1s to traverse a distance of 50 -100 µm.

To investigate whether protons could elicit the observed wound response, we visualised dynamic proton levels, i.e. pH, both in the medium and in the plant cell walls. To do so we used the recently reported CarboTag-OregonGreen (CT-OG), a FLIM-based apoplastic pH probe^36^ in combination with ablation (Fig. 3a). The results revealed a significant acidification of the medium within seconds upon laser ablation, with the pH of the medium decreasing from 5.5 to 4.5 (Fig. 3b), which was transient and recovered to its pre-ablation level within 60 seconds. Using FAST-FLIM, we captured the proton emission from the ablated cell as a rapidly traveling hemispherical wave that spreads in time from the wounding site (Movie S4).

**Figure 3:**
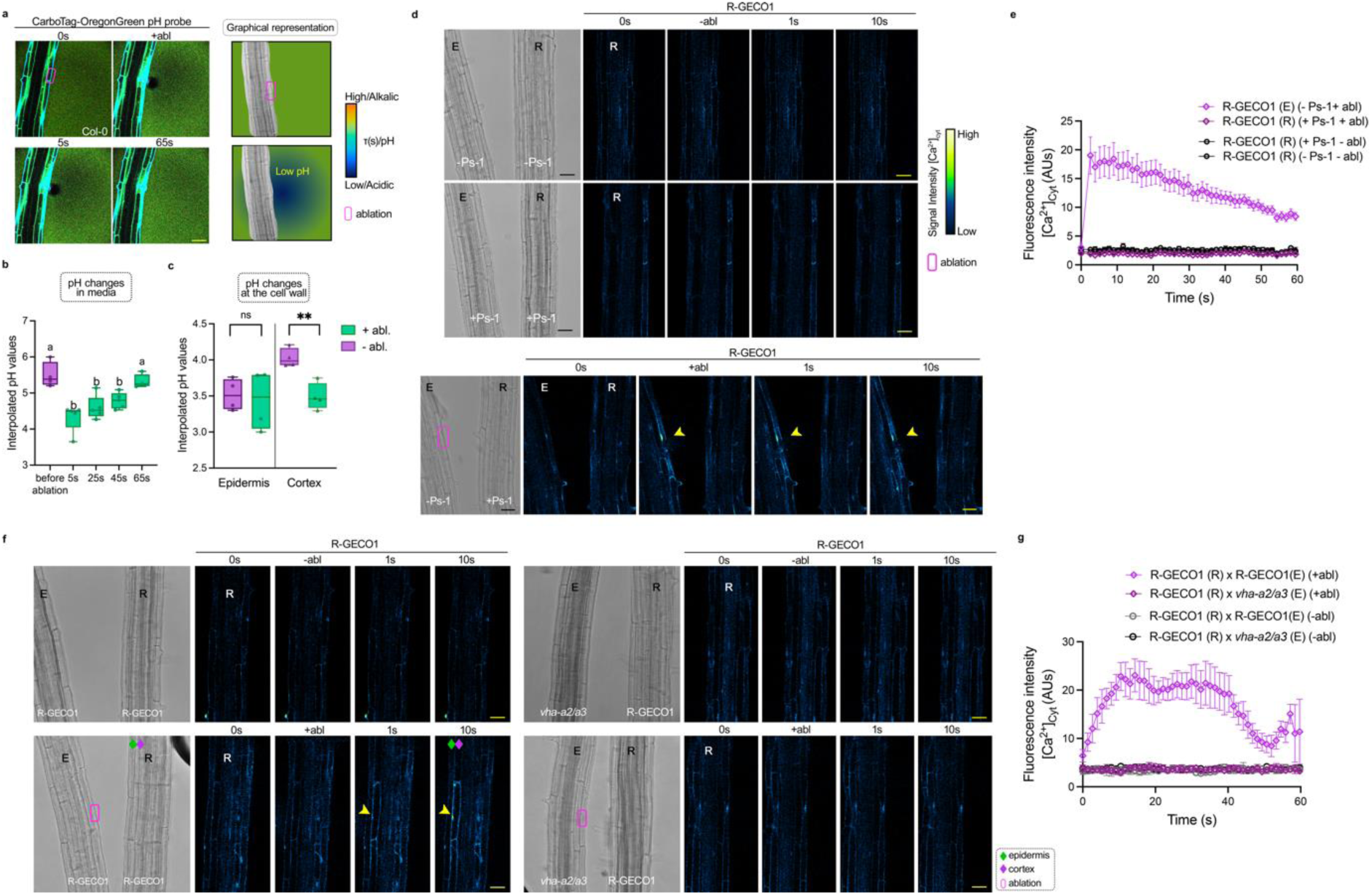
Rapid H^+^ released to environment upon wounding triggers a cortex-specific Ca^2+^ wave. a) Representative time-lapse images of five-days-old Col-0 seedlings treated with a CarboTag-OregonGreen FLIM-based pH probe (40 mM) to follow changes in pH at the cell wall before and after epidermal laser ablation. Graphical representation the lower lifetime values indicate an acidification of the media after wounding (pH values visualized according to the spectrum). b) Quantification of lifetime values transposed to interpolated pH values before and after ablation (n=5 with N=2, Paired two-tailed t-test, (a) is P < 0.05 and (b) P<0,005). The pH drops immediately in response to the ablation of an epidermal cell and recovers over a period of 65 seconds. c) Quantification of lifetime values transposed to interpolated pH values before and after wounding at the cell wall (n=5; N=2). The cortex/epidermis cell wall in receiver roots R experiences acidification in response to the laser ablation of epidermal cells in the emitter roots E. d) Seedlings expressing *At*R-GECO1 and treated with the proton pump inhibitor Protonstatin-1 (Ps-1, 20 μM) before and after ablation; there is no induction of the Ca^2+^ wave in the R compared to wounded control E roots. e) Quantification of [Ca^2+^]_cyt_ over time in the cortex of seedlings expressing *At*R-GECO1 that received or did not receive Ps-1 treatment (n=10, data represented as mean ± SD; N=3). f) Representative timelapse of *At*R-GECO1 receiver root R facing *At*R-GECO1 in control condition (E root) or *vha-a2/vha-*a3 double mutant (E root). Calcium wave induction R is observed only when facing *At*R-GECO1 emitter root. g) Quantification of [Ca^2+^]_cyt_ over time in the cortex of seedlings expressing *At*R-GECO1 facing *At*R-GECO1 or *vha-a2/vha-a3*, in AUs. The image in a) represents the results of two independent replicates while the images in d) represent the results from three independent replicates. a-d) scale bar, 50μm; green diamonds indicate the epidermal cell layer, while purple diamonds indicate the cortex. Magenta boxes indicate the ablation site and the yellow arrows the calcium wave. Yellow arrows indicate the calcium wave response. The signal intensity of the R-GECO1 calcium reporter (Green-Fire-Blue) is visualized according to the spectrum in 3d. AUs; Arbitrary Units.

The probe also gives access to estimate the pH changes in the cell wall of the plant. The cell wall pH in the R root decreased by half a pH unit in the cortex (Fig. 3c), i.e., an interpolated of pH 3.5, which correlates well with previously observations of a decrease in apoplastic pH (pH_apo_) in the cortex of Arabidopsis roots treated with an elicitor^37^. As a control, we tested the effect of the two-photon ablation laser on the probe itself; ablating the medium itself did not lead to a detectable change in the signal of the CarboTag-OG pH probe; this confirms that the observed rapid acidification is the result of wounding and not due to the use of a strong laser irradiation alone (Extended Data Fig. 4e, f). We confirmed these data by using the genetically encoded pH sensor SYP122-pHusion^37^; epidermal laser ablation also here resulted in a decrease in the relative pH in both the epidermal and cortical membranes (Extended Data Fig. 3 a, b).

These results paint a picture of protons as ultrafast signals that are emitted upon wounding. To investigate whether the protons are simply leaked from the damaged cell(s) or require proton pump activity which would suggest an active signal emission from E that these emitted protons are the signals perceived by the receiver R root to activate calcium signaling, we treated E and R or only R Col-0 seedlings with one of the proton pump inhibitors dicyclohexylcarbodiimidne^38^ (DCC-D; Extended Data Fig. 4a, b) or Protonstatin-1 (Ps-1; Fig. 3d; Extended Data Fig. 3c); in both cases, [Ca^2+^]cyt signaling was abolished (Extended Data Fig. 3f; Fig. 3e). Building on this, we investigated the double mutant *vha-a2/vha-a3* (referred to hereafter as *vha-a2/a3*; Fig. 3f) in the vacuolar ATPase subunit VHA-a2 and VHA-a3, which are known for their less acidic vacuole properties^39^. When *vha-a2/a3* roots were used as E, no [Ca2^+^]_cyt_ signal was induced in the R roots compared to the control (*At*R-GECO1) (Fig. 3g). To further confirm the role of vacuolar pH in wounding perception, we performed laser ablation experiments on both Col-0 and *vha-a2/a3* roots in media supplemented with fluorescein sodium (FS, 20 µM). In contrast to Col-0, where laser ablation induced acidification, *vha-a2/a3* showed a slight alkalinization of the environment (Extended Data Fig. 3e). Together, these findings demonstrate that H⁺ released from wounded roots serves as an ultra-fast messenger between plants (Extended Data Fig. 2). The absence of [Ca2^+^]_cyt_ signaling in the proton pump-inhibited and mutant roots underscores the critical role of active proton transport in plant communication.

### Rapid proton sensing following wounding induces a root avoidance mechanism

The obtained results raise the question regarding function has the rapid proton signaling in preventing damage in the receiver roots. To study this, we used an organic bioelectronic device, the Organic Electronic Ion Pump (OEIP)^40,41^ to selectively deliver H^+^ protons in close proximity of a root (Fig. 4a; Movie S5) without fluid flow. The OEIP is an electrophoretic (also denoted ‘iontronic’) delivery device that delivers ions and charged molecules from a source electrolyte to a target electrolyte via a charge selective membrane upon application of an electric field. The ions when they exit the device outlet they move solely by diffusion, in this way creating a local control of ionic concentration, that with a magnitude and range that scales with the applied electric field. First, the device was stabilised during an initial rest period of two minutes. The device was placed in a fresh agar solution for a 10-minute rest at 0 nA (depicted in Fig. 4 as ‘no H^+^ delivery – OEIP OFF’), followed by active delivery for 10 minutes at 400 nA (i.e. ‘H^+^ delivery – OEIP ON’) the electrophoretic delivery. Before testing whether there is a phenotypic effect on H^+^ treatment, R-GECO1 seedlings were subjected to five minutes of continuous H^+^ delivery (distance: 100 µm). The results show that continuous proton delivery is associated with elevated [Ca^2+^]_cyt_ in the cortex (Fig. 4a, b), with the concentrations demonstrating a pulsed pattern, which agrees with the ablation data (Fig. 4c, d; Movie S5).

**Figure 4:**
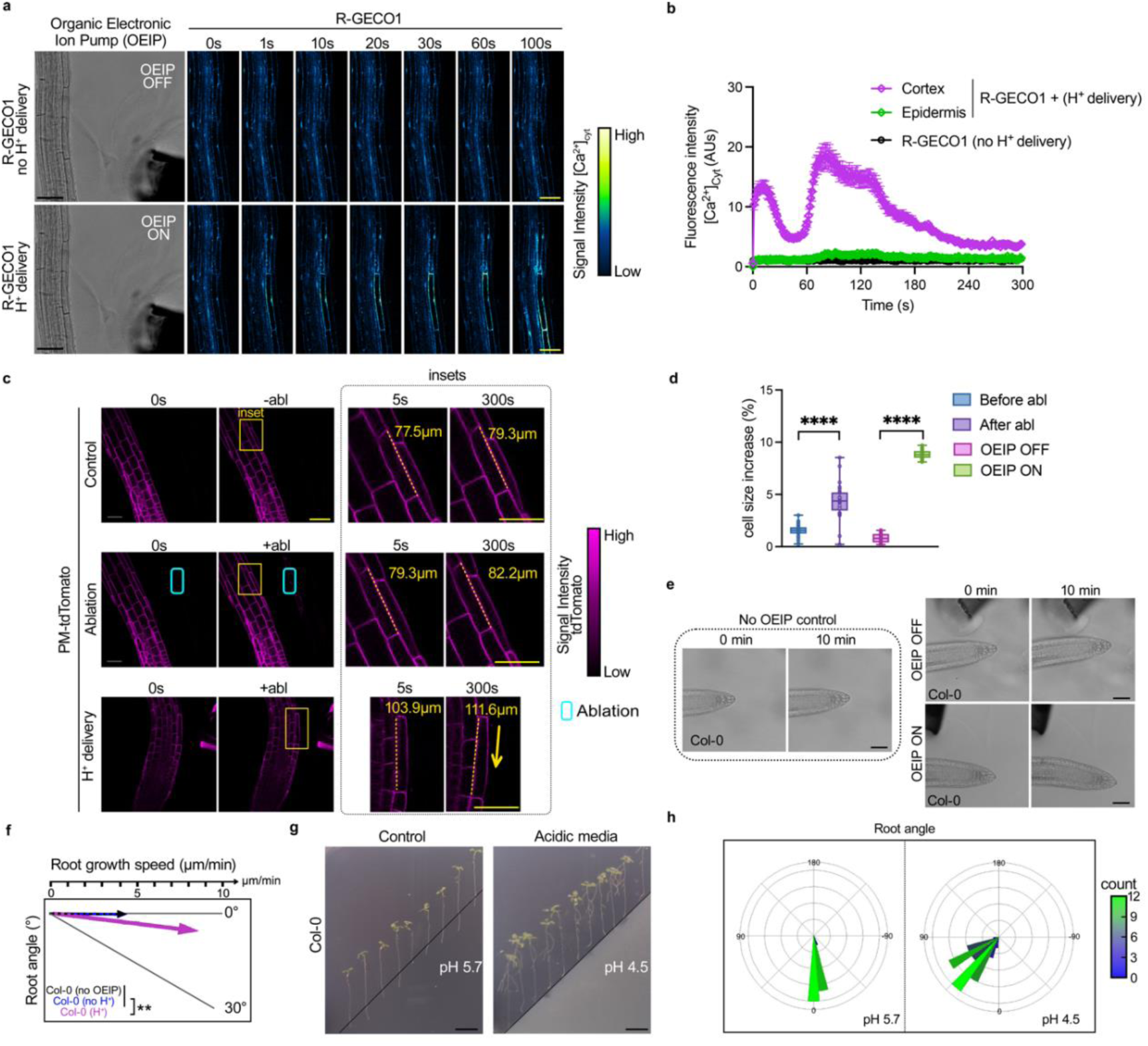
Root avoidance triggered by ultra-fast proton sensing. a) Representative time-lapse images of the Ca^2+^ wave in roots expressing AtR-GECO1 subjected to Organic Electronic Ion Pump (OEIP) delivery of H^+^ (400 nA). Panel (a) shows the immediate observation of cortex-specific Ca^2+^ signaling during local H^+^ delivery, while no Ca^2+^ signal is observed when the OEIP is turned off (control situation, white dashed lines (R)). Panel (b) presents the quantification of cytosolic calcium levels ([Ca^2+^]_cyt_) over time in the cortex and epidermis without and with H^+^ delivery by OEIP. c, d) Cell elongation in response to the H^+^ flux induced by laser ablation and OEIP. In both scenarios, cells exposed to H^+^ elongate significantly compared to the control. e) Brightfield images of Col-0 roots under control conditions without OEIP devices and with OEIP devices off (no H+ delivery) to assess basal root growth, as well as Col-0 with OEIP on for 10 minutes of continuous H^+^ delivery. Col-0 exhibits a growth rate that is 2-3 times faster. f) h) Images of seven-days old Col-0 seedlings transferred to media containing basal media (pH5.7) or acidic media (pH4.5) after 20h of growth. Seedling facing acidic media shows clear root avoidance comparing to basal media. i) Polar histograms are showing the root angles depending on the media facing the seedlings. Blue to Green scale represent the repartition of the root angle of n=20 seedlings. For panels d, e, data are represented as mean ± SD, n=10. The image in a, c, e, f, h are representative of 3 independent replicates. Statistical significance was determined by Student’s Paired t-test and represented as **p<0.01, ***p<0.001, ****p<0.0001. Scales for a, c, e, f are 50µm and 1 cm for g. The signal intensity of the R-GECO1 calcium reporter (Green-Fire-Blue) is visualized according to the spectrum in 4a. AUs; Arbitrary Units.

The results that wounding and OEIP-facilitated H^+^ diffusion are sensed by neighbouring roots led us to hypothesise that H^+^ fluxes should induce a Pattern Recognition Receptor (PRR), which is often implied in wound sensing and apoplastic pH changes^42,43,27^. PRR-PEPR1^27^ was selected as a candidate mutant to undergo single-cell ablation and proton treatment with OEIP (Extended Data Fig. 4c, d). The results revealed that PEPR1 is localised to the epidermis after 10 minutes of treatment. This implies that H^+^ fluxes might serve as a universal signal that triggers PRR-mediated responses, potentially broadening our understanding of how plants perceive and respond to environmental stress at the cellular level.

Next, we aimed to investigate the extent of acidification induced by both methods; OEIP-facilitated H^+^ diffusion and local acidification following single-cell laser ablation. We used MS media supplemented with 20 µM fluorescein-5-(and-6)-sulfonic acid, trisodium salt (FS)^44^ to quantify pH changes (Extended Data Fig. 5d-i). As a reference, MS plates supplemented with FS at pH 4-6.5 were prepared and imaged, with the fluorescence ratios quantified and displayed against log(pH) values; an R² of 0.999 validated the use of this approach for quantifying pH changes (Extended Data Fig. 5g). OEIP driven H^+^ delivery to the MS media supplemented with FS showed significant and rapid acidification (Extended Data Fig. 5d, e), with the pH falling to a minimum of approximately 3.5 (Extended Data Fig. 5h). Significant acidification was also observed following single-cell laser ablation (Extended Data Fig. 5f), with the pH decreasing to a value between 3.5-4.5 (Extended Data Fig. 5i). This validates the previously employed method and provides crucial information for subsequent assays of root avoidance; notably, the FLIM-based probe ablation experiment (Fig. 3a, b) indicated that the pH falls to 4 following wounding.

To further investigate acidification dynamics following OEIP H^+^ delivery, a digital representation of the experimental setup, including iontronic device and target environment, was implemented as a finite element model using COMSOL Multiphysics 6.2. The distribution of potential gradients and ionic concentration gradients were numerically solved using the Nernst-Planck equations while mimicking experimental conditions. The estimated change in concentration vs time caused can be seen in Extended data 5a-c. During the initial 10 min rest, the model indicates that the local pH close to the device is slightly below the surrounding media due to passive leakage. Fig 4a indicates however that the pH decrease due to passive leakage was however not sufficient to trigger a measurable response in elevated [Ca^2+^]_cyt_. During delivery, the model indicates that this OEIP-mediated H^+^ delivery generates a local pH3-3.6 within the region of 50-150 µm from the device outlet (Extended Data Fig. 5b, c), in agreement to experimental estimations using FS and fluorescent ratios (Extended Data Fig. 5e-i). See Supplementary materials for method details, parameter tables, initial values, and assumptions made in this analysis. The dose-dependent response is also in agreement with the wounding results (Fig. 1b-e), where the receiver roots responded to the wounding of the emitter only if the distance was short, i.e. sufficiently large drop in pH was needed to trigger response.

Next, we investigated the possible physiological role(s) of H^+^ driven root-to-root responses and how a reduction in vacuolar H^+^ levels stimulates cell elongation and root growth^45^ (Fig. 4c-f). The OEIP-mediated delivery of H^+^ to Col-0 root tips was associated with a significant increase in growth speed (2-3x faster) relative to observations from the control (Fig. 4e, f; Movie S6). Interestingly, analyses of root angle (Fig. 4f) during recordings of root growth (Fig. 4e, f) revealed that roots exposed to H^+^ grew faster and at a greater angle than roots that were not exposed to H^+^ (Fig 4g). This observation prompted us to further analyse root cell elongation on the side of the root exposed to H^+^. Both OEIP and laser ablation treatments resulted in a significant increase in cell elongation compared to observations from the control (Fig 4c, d). Root cell elongation on the side of a root exposed to H^+^ triggers roots to bend “away” from the acidic environment. To further demonstrate the induction of a root avoidance mechanism following H^+^ flux, we performed split agar experiments^46^. More specifically, Col-0 seedlings were placed on MS agar (pH 5.7), with some facing the MS media with a pH value of 5.7 (control) and others facing MS media with a pH value of 4.5 (acidic conditions) (Fig. 4g). Growth was recorded over 16h and the root angle was quantified (Fig. 4h). Seedlings exposed to acidic media showed significant root bending, with angles between 45 and 65 degrees. This physiological response supports our findings that H⁺ drives rapid below-ground plant reactions and aligns well with the acid growth theory, which suggests that an increase in H⁺ concentration in plant cells leads to the activation of cell wall-loosening enzymes, promoting cell expansion and growth^47^. Therefore, the acid growth theory provides a plausible explanation for the physiological responses observed in our findings work (Fig. 5).

**Figure 5:**
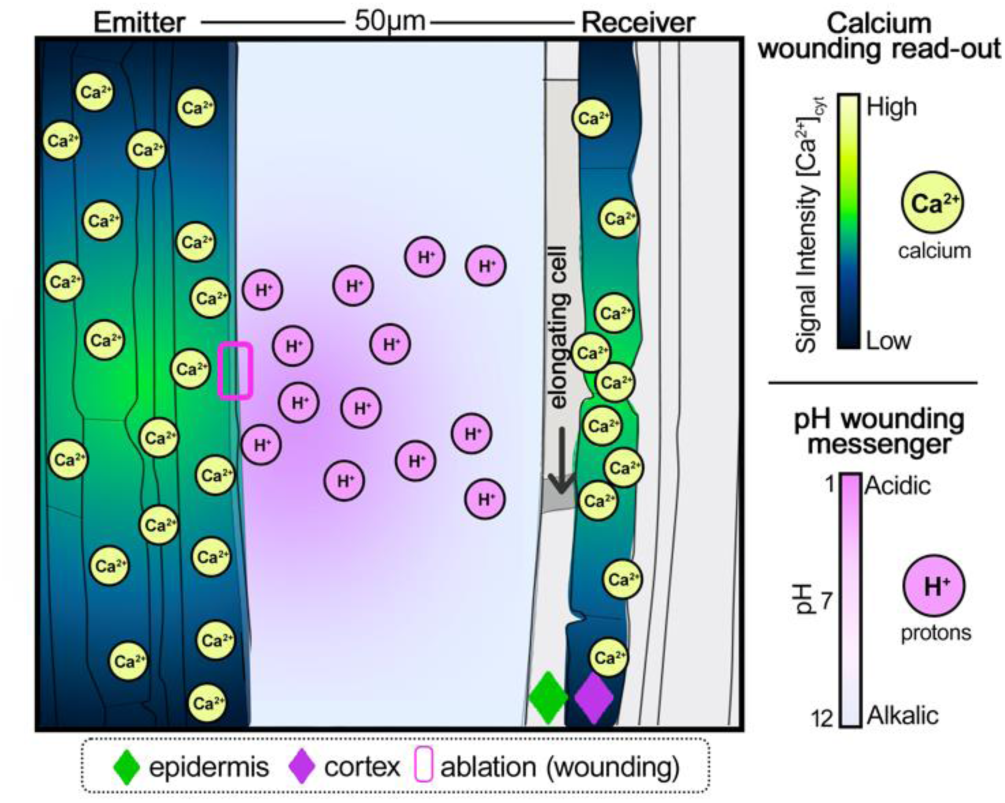
Model of root-to-root communication/signalling: ultra-fast proton release from damaged root induces cell elongation in neighbouring root. This model describes the wound induced root communication mechanism mediated by proton release. More specifically, the emitter root upon wounding (single cell laser ablation; indicated by a magenta box) releases protons, which acidify locally the surrounding environment and reach the receiver neighbouring root within seconds. This proton release initiates a cortex specific calcium wave response in the receiver-unwounded root, promoting cell elongation, which in turn results in a root avoidance phenotype. Hence, pH acts as the messenger of the wounding signal, whereas, the cytosolic calcium wave is used as the signal perception and read-out. Epidermis and cortex cell types are indicated by green and purple diamonds, respectively.

## Discussion

We visualised and confirmed rapid proton diffusion during root wounding (Fig. 3, 4) which was perceived by a neighbouring root (Figs. 1, 2). The rapid release of cell content during wounding induces the significant release of H^+^, which can acidify the root environment by up to 1 pH unit (Fig. 3b, Extended data Fig. 5g-i). This results in a rapid and robust non-canonical response from the plant in the form of increased cell elongation, along with a root avoidance mechanism, within five minutes of wounding (Fig. 4c-h). Recent studies have described how H^+^ levels have a pivotal role in plant development; more specifically, an acidic apoplastic pH (pH_apo_) promotes root growth while an alkaline pH_apo_ increases cell wall stiffness to boost plant defense and protection against osmotic stresses^48, 27^. Here, we also provide new evidence that rapid H^+^ diffusion serves as a root-to-root communication mechanism, which consolidates the fact that local H^+^ fluxes represent a direct link between external stimuli present in the root environment (Extended Data Fig. 2b) and plant physiology. The presented research also supports the acid growth theory, which is already strongly accepted among the scientific community and describes the link between auxin, pH, and cellular expansion^47^. Previous research reported that – within the hypocotyl and root elongation zones – the kinase TMK1 is responsible for the H^+^-ATPase phosphorylation-induced pH_apo_ acidification, auxin signaling, and thus, cell expansion^49,50^, while another recent study revealed that danger-associated peptide Pep1 induces the hyper-expansion of cells within the transition zone (TZ) upon the alkalinisation of pH_apo_ and auxin accumulation in the TZ^51^.

Another interesting feature of this rapid proton perception is the elevated [Ca^2+^]_cyt_ in the cortical cell file (Figs. 1, 2, 3 and 4), as well as the significant acidification at the cell wall (Fig. 3c; Extended Data Fig. 3a, b). It was reported that pHapo acidification in the cortical cell file can modify the internal membrane pH while the cytosol of other root cells maintains a stable pH^37^. Notably, the fact that cortical cells can have a slightly different pH than other root tissues suggest tight regulation of the AHA and thus, the Proton Motive Force (PMF). According to prior findings, pH changes in cortical cells could affect essential cellular processes such as microtubule maintenance, along with the expression of transporter or cell-surface receptors that are crucial for the maintenance of cell wall integrity and the perception of damage^43,52^. This aspect should be further investigated to better understand the tight regulation of H^+^ fluxes in root cells as well as the mechanisms underlying proton sensing. Overall, our data propose a novel form of ultra-fast root-to-root communication based on rapid H^+^ diffusion as a consequence of wounding (Fig. 5).

## Methods

### Growth conditions and plant material

The *A. thaliana* seedlings used in all of the experiments were six-days-old, and grown vertically in an *in vitro* set-up (16h of light at 150 μmol photons m^−2^ s^−1^, 62% RH, 22°C day, 18°C night) on ½ MS media (Murashige and Skoog Basal salt mixture including vitamins, MES buffer adjusted with KOH to pH 5.7, 0.8% Plant agar; all from Duchefa Biochimie, Haarlem, Netherlands). Seeds were surface sterilised with 70% ethanol for 10 min, rinsed with sterile water, sawn with 0.1% agarose and stratified for 1 day at 4°C in the dark. For seed propagation, plants were placed in a long-day growth room (16 h light at 150 μmol photons m^−2^ s^−1^ and 22°C, 8 h dark at 18°C) in a pot with a specific soil mixture (1:3 mixture of Agra-vermiculite + “yrkeskvalité K-JORD/krukjord” provided by RHP (MG ‘s-Gravenzande) and Hasselfors garden (Örebro, Sweden), respectively).

Seeds used in the study were *A. thaliana* R-GECO1^29^, and GCaMP3^30^ for [Ca^2+^]_cyt_ visualisation; PEPR1::PEPR1::GFP in *pepr1pepr2* ^53^ were used in the PEPR1 localisation experiments; while SYP122-pHusion^37^ served as ratiometric pH sensors; *vha-a2* x *vha-a3 A. thaliana* double mutant plants in subunit C of the vacuolar H^+^-ATPase isoform A2 and A3, were used to investigate the regulation of proton fluxes^54^. For the cell elongation measurements, pUBQ10::RCI2A-tdTomato was used^55^.

### Pharmacological seedling treatment

The six-days-old R-GECO1 seedlings were treated with a specific drug or without any additional substance (liquid ½ MS). The seedlings receiving proton pump inhibitors were treated with 50µM of dicyclohexylcarbodiimidne (DCCD, Sigma Aldrich, St. Louis, MO; D80002-25G) and 5µM of Protonstatin-1 (Ps-1, BiorByt, Cambridge, UK; orb1737682). Seedlings that received treatment were incubated with the proton pump inhibitors diluted in ½ MS in a micro-well plate in the dark and in *in vitro* chambers with agitation. For imaging, seedlings were placed next to each other in sterilized /pre-humidified glass chambers (Nunc Lab-Tek, Thermo Fisher Scientific, Waltham, MA) with ½ MS (1% plant agar) on top of the root to keep it humid. A single cell of the epidermal cell file of the remitter R-GECO1 root (E) was ablated with a laser and calcium wave induction in the receiver root (R) was recorded for 60 s at 1 frame per second (detailed method provided in the section about calcium signal analysis). In the case of control plants, roots were incubated in ½ MS without any proton pump inhibitor in the dark and with agitation; laser ablation proceeded in the same manner. Each dataset comprises 10 technical replicates for each of the three biological replicates.

### Proton delivery via Organic Electronic Ion Pump

The OEIP devices were fabricated according to previously reported procedures**^56^**. Briefly, in the current work a cation-exchange membrane based on 2-Acrylamido-2-methyl-1-propanesulfonic acid sodium salt (AMPS) crosslinked with polyethylene glycols diacrylate (PEGDA) was placed in the glass capillaries of smaller hollow (25 µm inner diameter) with photoexposure time of 170 min. The OEIP reservoir was filled with 0.01M HCl (Fig. S4a). In the experiments that involved H^+^ delivery via OEIP, six-days-old R-GECO1 seedlings were used to follow calcium signaling, while Col-0 wild-type plants were used for root growth measurements. Individual roots were placed in a pre-humidified round glass chamber (Nunc Lab-Tek.) with ½ MS (1% plant agar) placed on top of the root to maintain humidity. First, the device was stabilised during an initial rest period of two minutes (Fig. S4a-b). The OEIP was placed near the root with a micromanipulator (Sensapex uM-TSC, zero drift) and a current of 400 nA was applied. Control videos of each root were recorded for 10 min before treatment, with 1 frame per 0.73 seconds. The same recording parameters were used during proton delivery. Each data set includes 10 technical replicates per experimental condition for each of the three biological replicates (n=10 for N=3). Methods for the H+ delivery simulation is fully described in the Supplementary information.

### Confocal imaging and real-time analysis

Seedlings were placed into a pre-humidified sterile glass chamber (Nunc Lab-Tek)^31,32^, and covered by either a piece of ½ MS medium with MES buffer (imaging) or a piece of ½ MS medium without MES buffer (pH analysis). The glass chamber was then closed with a lid to maintain the humidity necessary for *in planta* imaging. For confocal imaging, A laser scanning Leica Stellaris 8 microscope (Leica, Wetzlar, Germany) fitted with a x40 glycerol objective was used for confocal imaging. The fluorescence signals for mRFP (excitation 560 nm, emission 588-658nm), GFP (excitation 488 nm, emission 500-550 nm) were detected with a HyD hybrid photodetector (scanning speed 400 Hz). The SYP122-pHusion fluorescence signal was detected at 488 nm / 405 nm (emission at 500-580 nm), and the 488/405 ratio was calculated as described in Kresten et al., 2019 with an imaging time of 15 min (with 1 frame per second to yield an average of 900 frames).

### Calcium signal analysis

Increases in cortical and epidermal [Ca^2+^]_cyt_ were monitored in the epidermal and cortical cells of the root neighbouring the laser ablated root. When imaging the Ca^2+^ wave in UBQ10pro::GCaMP3 and R-GECO1 intensiometric Ca^2+^ reporter lines, the laser power was set to 8%. Images were recorded every second (s) for 60 seconds to yield 60 images (Fig. 1-4, Supp. Movie S1-S3 prior to ablation. For the UBQ10pro::GCaMP3 reporter lines, we used the previously described GFP settings (excitation 488 nm, emission 500-530 nm), while the settings reported by Keinath et al. (2015) were used for R-GECO1 reporter lines (excitation 561 nm, emission 620-650nm). The fluorescence intensity of the region of interest (ROI) (neighbouring undamaged cell 50-150µm from the ablated root at the beginning of calcium wave induction) was measured using LasX software (version 2.0.0 14.332). We obtained measurements from 10 independent roots (technical replicates, n=10). Because we used ten technical replicates for each three biological replicates (N=3), there were 3 x (60 x 10) frame, or a total of 900, pictures illustrating the relative fluorescence intensity of [Ca^2+^]_cyt_ for each data set in arbitrary units (AUs). We consider this to represent relative fluorescence intensity of [Ca^2+^]_cyt_. The x=0 s timepoint correspond to the fluorescence intensity measured before ablation and x=1 s to fluorescence intensity measured at 1s after laser ablation. Movies were generated using the LasX software provided with Stellaris 8 DIVE (Version 2.0.0 14.332, copyright 1997, 2015, Leica). To measure calcium signaling in various root zones, the emitter root was ablated in MZ and calcium waves were recorded in different zones of the receiving root (DZ, EZ, RAM; Fig. 2A-B).

### Laser ablation

For laser ablation, a STELLARIS 8 Multiphoton/Confocal Microscope (Leica DMi8), equipped with a x40 glycerol objective, was coupled to Mai Tai Multiphoton laser (Spectra Physics, Milpitas, CA). The 800 nm laser was used in FRAP mode, with 50-60% laser power with the MPF SP667 filter for epidermal ablation while scanning in (xyt). The cell walls of epidermal cells from six-days-old seedlings in the sterile glass chamber were subjected to triangular selection of the ROI before ablation. The pre-ablation time was set to 2 seconds, the ablation time was set to 2 seconds (necessary to burst the cell open), and one frame per second was recorded for the following 60 seconds. At least 10 ablation events (technical replicates, n=10) were performed on 10 pair roots per data set, each of which included three biological replicates (N=3), if less indicated in the figure legend.

### FLIM imaging and analysis

FLIM images were recorded using a Leica SP8 multiphoton system with a pulsed Coherent Chameleon Ti:sapphire laser. Oregon green CarboTag was excited with a 980 nm line. Imaging was performed with a 40× water immersion objective and images were recorded at 512x512 pixel resolution, at a line scanning speed of 400 Hz. Emission was collected using a spectral window of 50 nm bandwidth centred on 525 nm onto a Leica HyD SMD hybrid photodetector. After image acquisition, Leica FLIM software was used to select and bin ROIs. A one-component tail fit (2-12ns) was used to determine the fluorescence lifetime. A decreased lifetime reflects an acidification whereas an increased lifetime value reflects an alkalinization. For this experiment, 5 technical (n=5) replicates were used per data set on 5 individual roots, each of which include 2 biologicals replicates (N=2) for the controls (measure in the media) and 5 pair of roots (measure in media and at CW; technical replicates, n=5) with N=2.

### pH analysis

For the pH changes induced by H+ delivery and laser ablation, 20μM of Fluorescein-5-(and-6)-Sulfonic Acid trisodium salt (FS) were added to the ½ MS (pH adjusted with KOH at 5.7, without MES buffer). The preparation and imaging were done as described in NBC Serre et al., 2023^44^. Images in Extended Data 3 are displaying the 488/405 nm (Green/Magenta) ratio, where green pixels depict pH values between 4-7 (acidic) and magenta pixels depict pH values between 7-9 (basic). For the reference, the 488/405 nm ratio was calculated after imaging of ½ MS (supplemented with 20μM FS) with different pH values from pH4 to pH5. Fluorescence intensities were plotted after logarithmic transformation (log(pH)) and third-degree polynomial curve was obtained with an r^2^ of 0, 999, validating the robustness of the method. For the pH change in the media (with H^+^ delivery or ablation), 10 square ROIs were analysed per picture, over 10 technical replicates (n=10), with respectively 3 biological replicates (N=3).

### Statistical information and software used

All experiments involved three biological replicates (N=3), each of which was subjected to 10 technical replications (n=10) except for the FLIM-based probe were 5 technical replicates (n=5) were used. Unpaired student’s t-tests was performed to determine the statistical significance of the results returned by technical replicates (n=10) and biological replicates (N=3), respectively; these statistical analyses were performed in Prism GraphPad (GraphPad Software, La Jolla, CA). The measurements were obtained using Fiji, Image J (National Institute of Health, http://rsb.info.nih.gov/ii). The pH interpolation analyses were performed in R Studio (script available in suppl. information SI2) for Fig. 3. The polar histogram representing the root angles displayed in Fig. 4i were obtained with the R Studio script available at SI3.

### Author contributions

PM initiated the project, while JD planned, designed, and conceived the project with input from PM. JD and PM wrote the manuscript with input from all co-authors. JD performed the ablation experiments, the root avoidance essay along with the data analysis. LT and JD performed the OEIP experiments, under the guidance of IBW, TAS, TN and ES. NZ helped with Gluconobacter while SJ with pH analyses. IBW fabricated OEIP devices, TAS performed finite element modelling of the OEIP delivery. LMF did the ablation experiment and cell elongation quantification along with the OEIP experiment with LT. AM performed laser ablation experimnets, data analysis and figure correction with JD. BM and JD did the FAST-FLIM under the guidance of JS.

## Acknowledgements

We would like to thank Åsa Strand, Charles Melnyk, Rishikesh Bhalerao, Steffen Vanneste and Matyáš Fendrych for their valuable comments on the manuscript. Furthermore, we thank to Laura Bacete, Eugenia Russinova, Karin Schumacher and Clara Sanchéz-Rodriguez for sharing published material. We acknowledge the Plant Growth and Microscopy facilities at UPSC for their assistance with the research.

## Data availability

All data and materials are available in the main text or the supplementary materials.

---

**Extended Data Figure 1:**
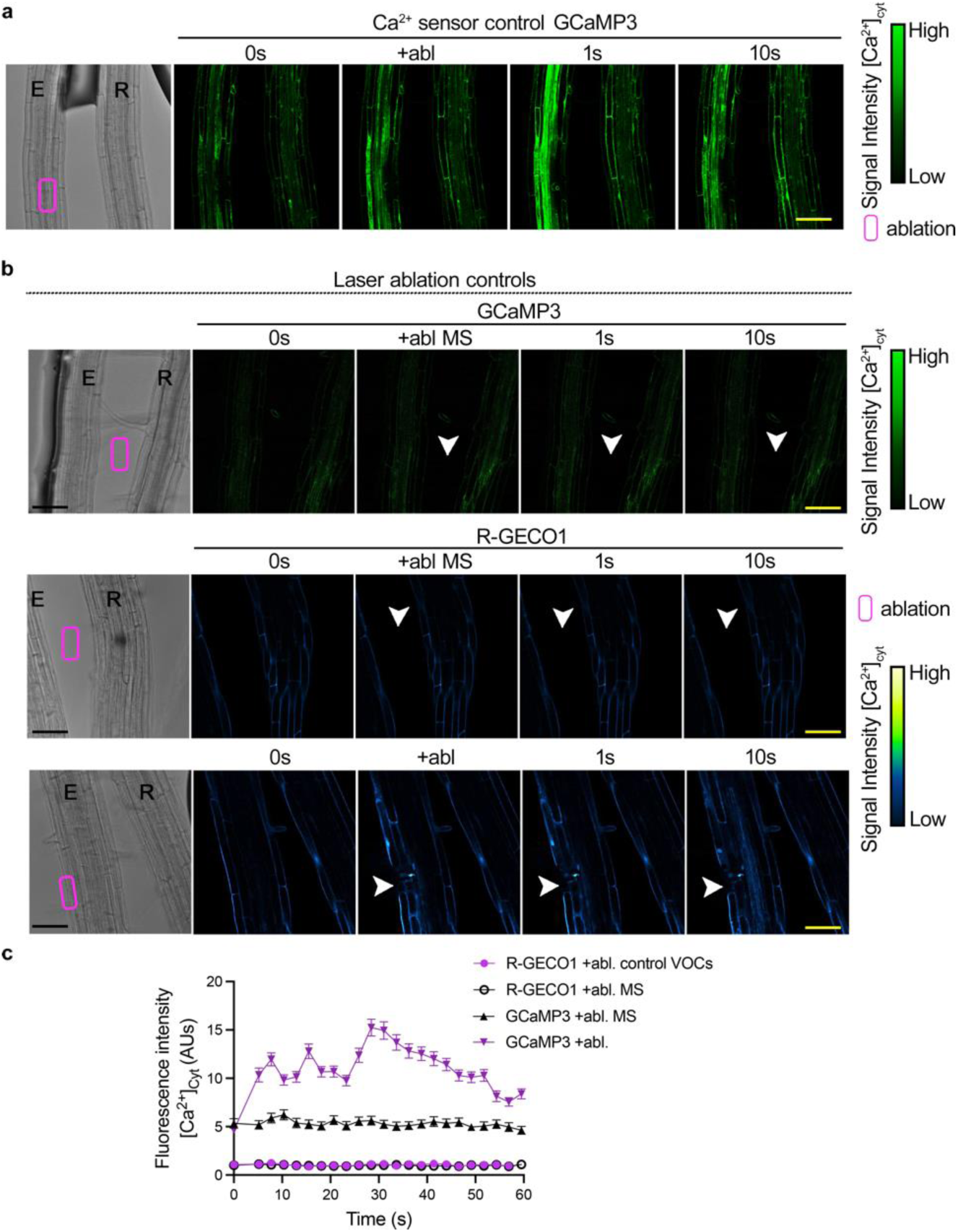
Using Ca^2+^ waves to visualise the perception of signals emitted by wounded plant cells. a) Representative time-lapse imaging of the AtUBI10::GCaMP3 tensiometric calcium reporter line (green) over 60 seconds after the ablation of the epidermal emitter E root (magenta boxes). Ablation induces an elevation in cytosolic Ca^2+^ ([Ca^2+^]_cyt_) in the cortex of the receiver root R and throughout all tissues of the emitter root E. b, c) Control experiments to assess the effect of the two-photon laser, which does not cause a Ca^2+^ wave when applied to the media surrounding the root. Ablation was performed b) in the media between two roots of AtUBI10::GCamP3 and *At*R-GECO1, and c) on an epidermal cell facing away from the neighbouring root, with neither showing an induction of Ca^2+^ signaling. d) Quantification of cortex-specific fluorescence intensity of cytosolic calcium ([Ca^2+^]cyt) from (a-c) (n=10 for each condition). Data are represented as mean ± SD. The images in (a-c) represent the results from three independent replicates (N=3). Scale bars for (a-c) are 50 μm. Magenta boxes indicate the ablation region of interest (ROI), and white arrows point to the ablation site. The signal intensity of the R-GECO1 calcium reporter (Green-Fire-Blue) is visualized according to the spectrum in 3d. AUs; Arbitrary Units.

**Extended Data Figure 2:**
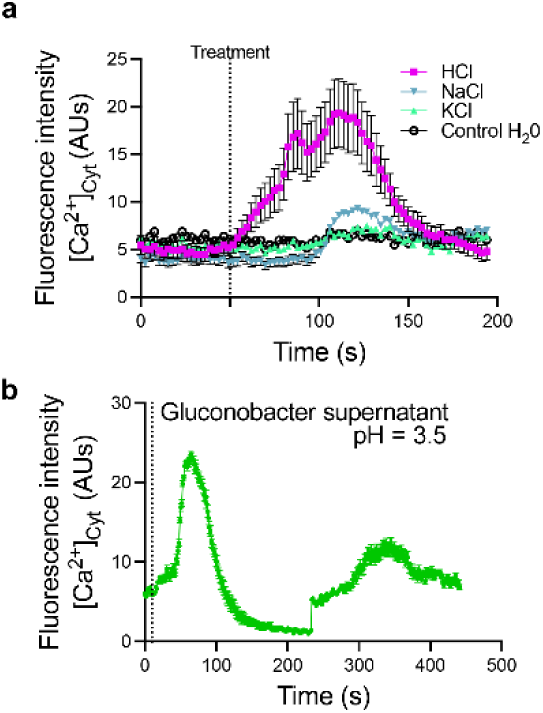
H^+^ sparks a cortex Ca^2+^ wave in unwounded roots. a) Graph representing the Ca^2+^ wave induced in *At*R-GECO1 roots over three minutes after inserting the root tip in KCl, NaCl, HCl respectively at 1mM and H_2_0 as control at t=50s. b) Quantification of the cortex Ca^2+^ wave in seedlings expressing AtRGECO1 after the application of *Gluconobacter oxydans* supernatant, which is rich in organic acids and shows a pH of 2.5 at t=15s. (a-b) Both treatments shows calcium induction in the cortex. The data represent the mean ± SD, with respectively n=10 and n=7, for 2-independent replicates(N=2).

**Extended Data Table 1:** Table displaying the diffusion coefficients and timescales calculated for different classes of signals as a function of root-to-root surface distances over two dimensions. Protons are shown to be the fastest at traversing distances.

| | *D* (m<sup>2</sup>/s) | $\tau_D$ @ 50 μm | $\tau_D$ @ 100 μm | $\tau_D$ @ 250 μm | $\tau_D$ @ 500 μm | $\tau_D$ @ 1 mm |
| --- | --- | --- | --- | --- | --- | --- |
| Proton (H<sup>+</sup>) | 9.3 × 10<sup>-9</sup> | 0.3 s | 1 s | 7s | 27 s | 108 s |
| Small ion (Na<sup>+</sup>) | 1.3 × 10<sup>-9</sup> | 2 s | 8 s | 47 s | 188 s | 751 s |
| Hormone (IAA) | 6.7 × 10<sup>-10</sup> | 4 s | 15 s | 94 s | 375 s | 1 500 s |
| Protein (GFP) | 8.7 × 10<sup>-11</sup> | 28 s | 111 s | 694 s | 2 777 s | 11 111 s |

**Extended Data Figure 3:**
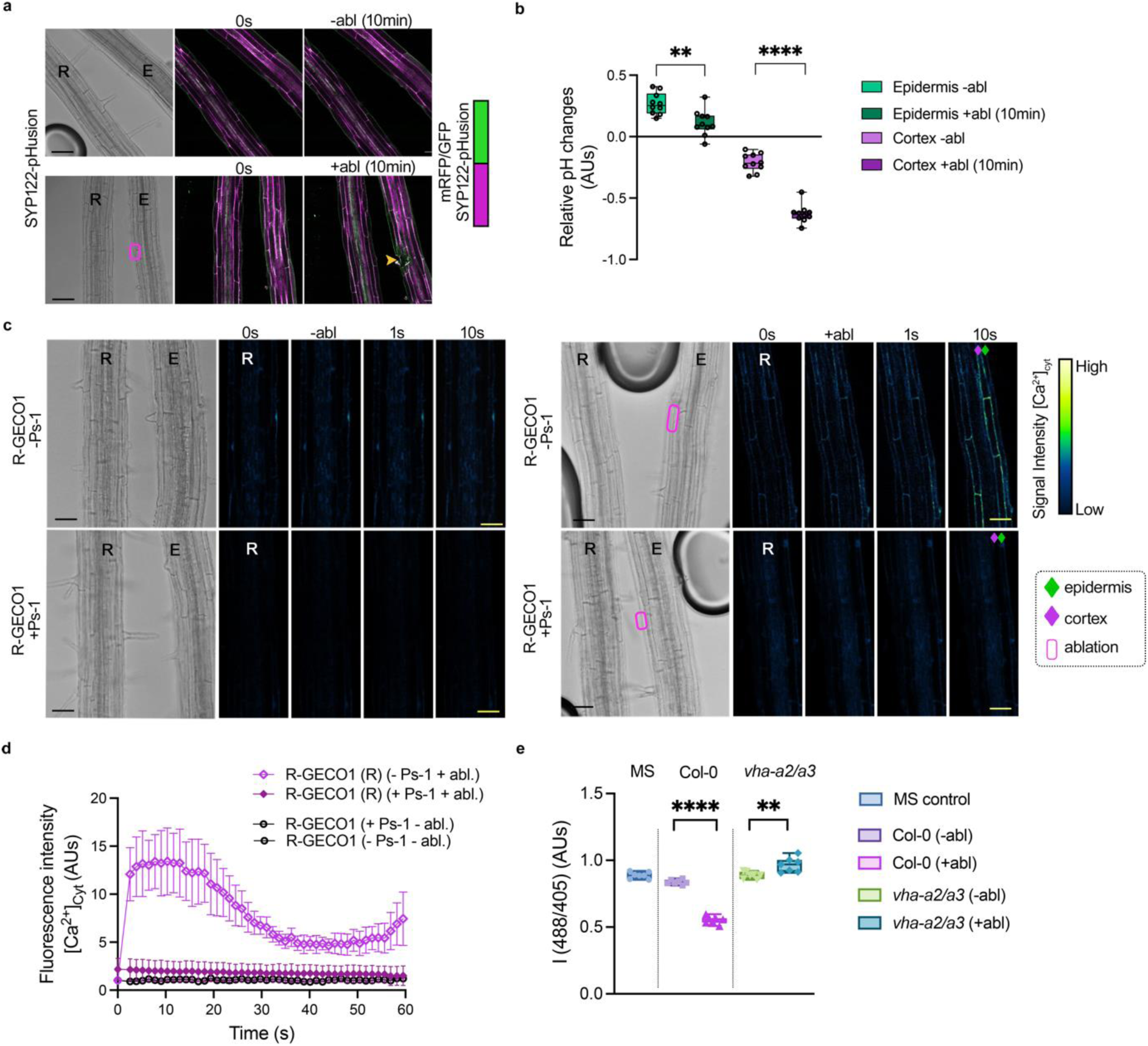
Acidic pH is perceived at the cortex-epidermis cell wall. a) Time-lapse imaging of the genetically encoded pH sensor SYP122-pHusion was recorded before and 10 minutes after epidermal ablation, with ratiometric images obtained (Green/Magenta). b) Quantification from (a). The relative pH changes were calculated in arbitrary units (GFP/mRFP) and revealed a significant acidification of epidermis and cortex cell walls in R after single-cell laser ablation. Data are represented as mean ± SD, paired Student’s t-test of n=10, **p<0.001, ****p<0.0001. c) Seedlings expressing *At*R-GECO1 and treated with the proton pump inhibitor Protonstatin-1 (Ps-1, 20 μM) before and after ablation (magenta boxes); there is no induction of the Ca^2+^ wave compared to control roots where cortex specific Ca^2+^ signalling can be observed. Quantification of [Ca^2+^]_cyt_ over time in the cortex of seedlings expressing *At*R-GECO1 that received or did not receive Ps-1 treatment (n=10, data represented as mean ± SD). Fluorescence images displayed R root from brightfield image. e) Box plot representing the ratiometric intensity (488 nm / 405 nm) obtained with 20μM Fluorescein-5-(and-6)-Sulfonic Acid trisodium salt (FS) measure in media in control conditions, and after laser ablation of Col-0 root and double mutant *vha-a2/a3* root. Ablation in *vha-a2/a3* shows alkalinization of the media whereas Col-0 laser ablation induces significant acidification of the media. Data are represented as median ± SD, unpaired student’s t-test of n=7 with **p<0.001, ****p<0.0001. The signal intensity of the R-GECO1 calcium reporter (Green-Fire-Blue) is visualized according to the spectrum in c. AUs; Arbitrary Units.

**Extended Data Figure 4:**
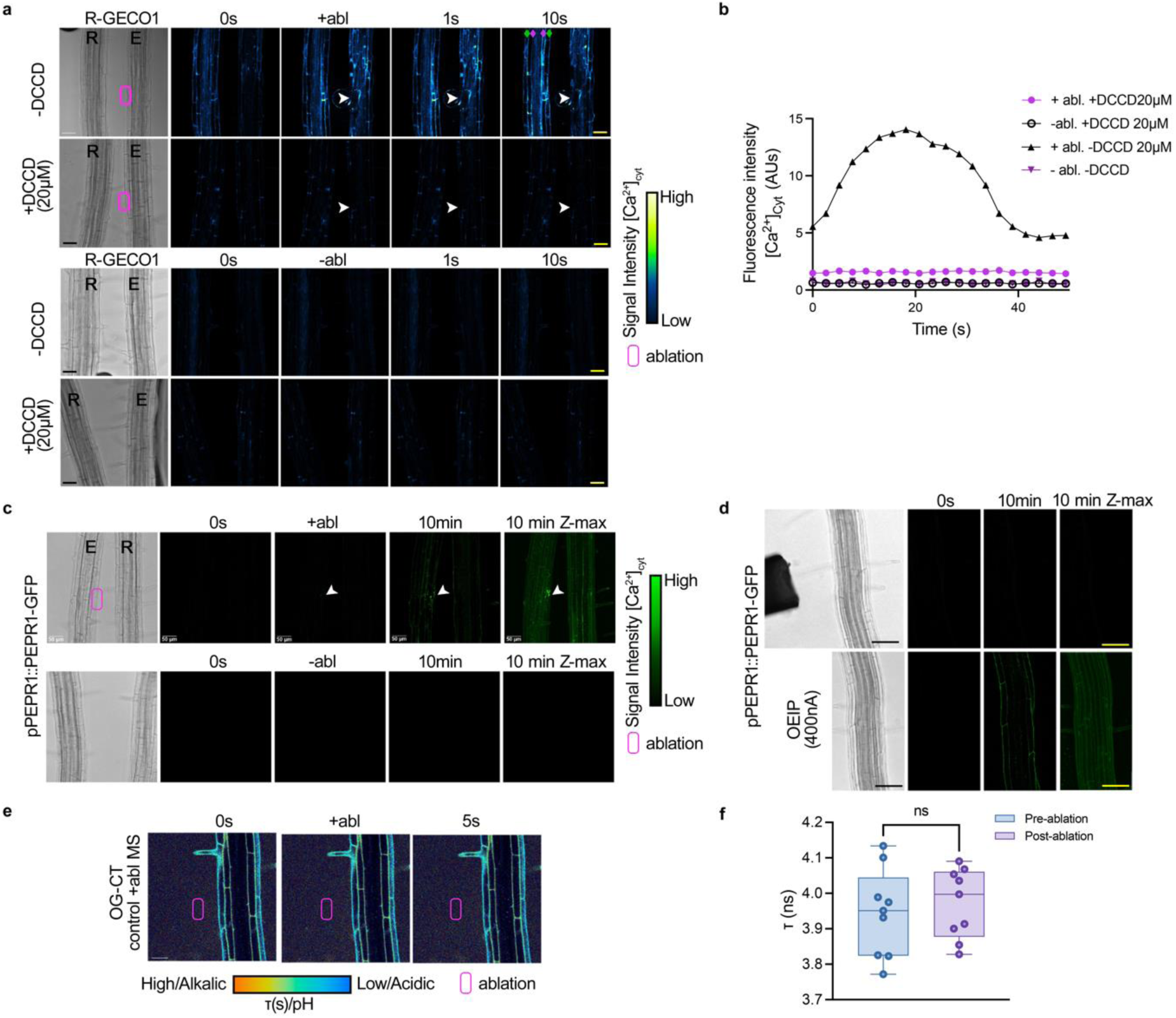
Acidic pH is perceived at the cortex-epidermis cell wall. a) Five-day-old seedlings expressing *At*R-GECO1 were treated with a dicyclohexylcarbodiimide proton pump inhibitor (DCC-D, 20μM) for 30 minutes before and after epidermal ablation (red arrow), and showed no elevation of cytosolic calcium levels ([Ca2+ cyt) compared to control roots, where cortex-specific Ca^2+^ wave is typically observed. Quantification from (a) is demonstrated in (b) as mean ± SD. c, d) Time-lapse imaging of the PEPR1 reporter line (pPEPR1::PEPR1::GFP; Green) over 10 minutes before and after epidermal ablation, and (g) prior to Organic Electronic Ion Pump (OEIP) delivery of H^+^ (d). PEPR1 localisation was examined 10 minutes after ablation and revealed rapid localisation of the receptor in both emitter E and receiver R cells, and (d) after 10 minutes in response to H^+^ from OEIP. e) Five-day-old Col-0 roots treated with the FLIM-based pH probe OregonGreen-CarboTag (40μM) were used as a laser control. MS ablation (magenta rectangle) was performed, and lifetime values showed no fluctuation in the media (f). Data are represented as median ± SD, paired Student’s t-test of n=10 (n=5 x N=2) with ^ns^p<0.01. The images in a, c and d represent the results of three independent biological replicates (N=3). Images in e are representing the results of five technical replicates with three independent biological replicates (n=5; N=3). Scale bars for (a, c and d) are 50μm; magenta zone illustrate the ablation zone, the white arrow the ablation site. The magenta diamond indicates the cortex cell layer while the green diamond indicates the epidermis cell layer. White arrows indicate the ablation area. The signal intensity of the R-GECO1 calcium reporter (Green-Fire-Blue) is visualized according to the spectrum in a and of the PEPR1-GFP signal in c. AUs; Arbitrary Units.

**Extended data 5:**
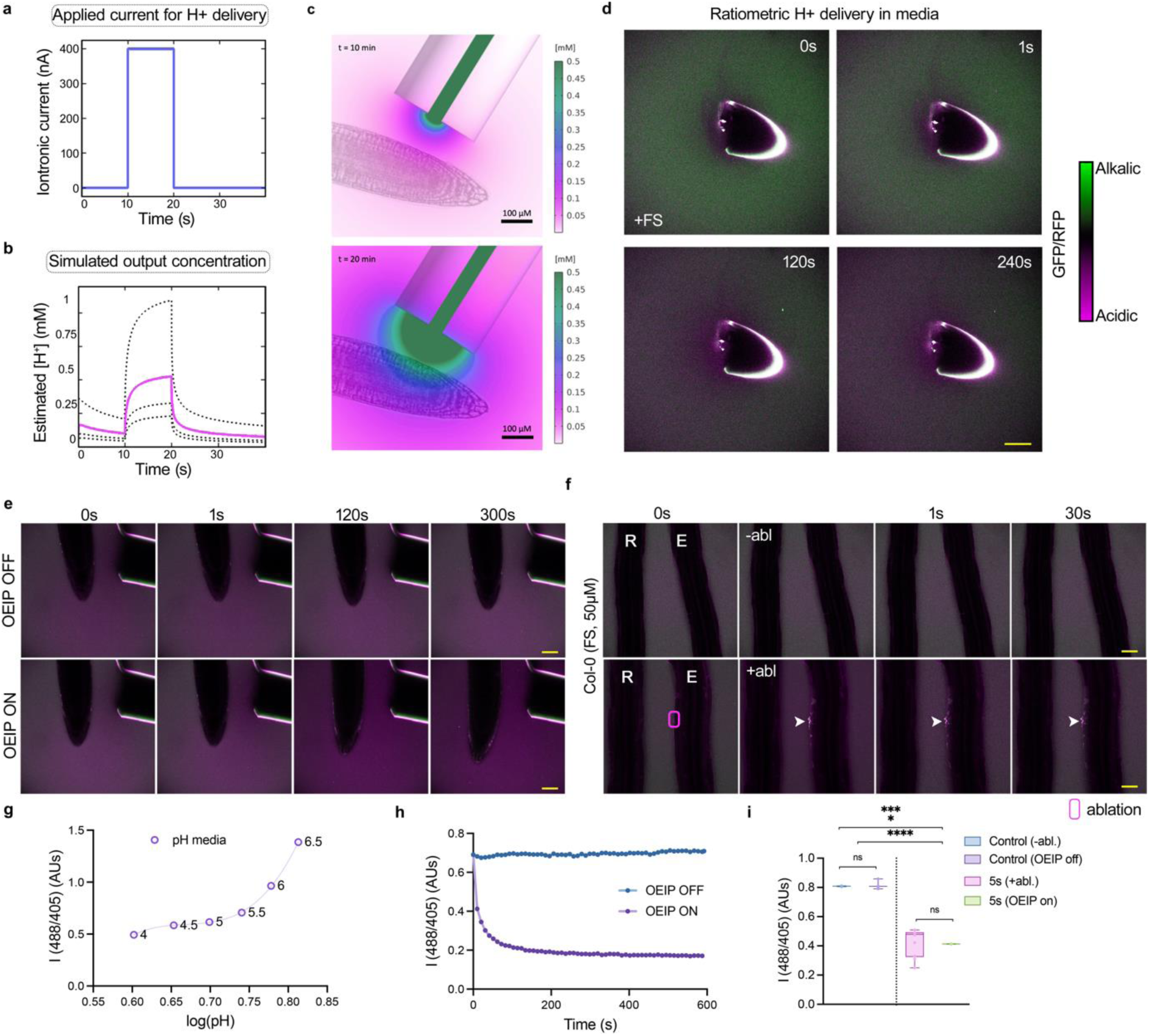
Acidification at the cell wall following OEIP treatment and epidermis ablation. a) Graph of the current (iontronic current in nA) applied over time with OEIP for proton delivery. b) Graph displaying the simulated proton output (mM) for a given distance of 50-200μm, red line indicating distance at 100 µm. c) Overlay representation of the estimated proton concentration (mM) during delivery with OEIP (before applied iontronic current (t = 10 min)) and at maximum concentration elevation (t = 20 min)). d) Time-lapse images of ratiometric H+ delivery in media supplemented with 20μM Fluorescein-5-(and-6)-Sulfonic Acid trisodium salt (FS). The media was found to turn acidic over four minutes of proton delivery. e) Images of Col-0 roots over a time interval of five minutes, without and with continuous OEIP-facilitated H^+^ delivery, in media supplemented with 20μM FS. f) Time-lapse images of Col-0 seedlings in media supplemented with 20μM FS before and after epidermal ablation in the emitter root E. After epidermal ablation, the media undergoes acidification. g) Graph displaying the 488/405 nm ratio in media (supplemented with 20μM FS) with different pH values. Fluorescence intensities are plotted after logarithmic transformation (log(pH)) to control the quality of the reflected pH values and the FS intensity measures. h) Graph of the 488/405 nm intensity ratios in media before and after H^+^ treatment on Col-0 root. The ratio decreases over time, with the pH decreasing during H^+^ treatment. i) Plot representing the 488/405 nm ratio measured in the media before and 5 s after ablation or H^+^ treatment (OEIP on); the results show that both approaches cause significant acidification in the media. (g-i) Data represent the mean ± SD where (g) includes five individual replicates, and (g, i) include three individual replicates. The images in (d) include two individual replicates (N=2), while the images in (e, f) include three individual replicates (N=). The scale bar for (d) is 200μm and 50 μm for (e-f). Magenta boxes indicate the ablation region of interest (ROI), and magenta rectangle highlight the ablation site. Statistical significance was determined by Multiple Student’s Paired t-test and represented as ****p<0.0001.

## Movies

**Movie S1:** Single-cell laser ablation in an emitter R-GECO1 epidermal cell facing a receiver R-GECO1 root, at 50µm distance.

**Movie S2:** Single-cell laser ablation in an emitter root (*S. lycopersicum*.) facing a receiver R-GECO1 root.

**Movie S3:** Laser ablation in media close to a receiver R-GECO1 root (R).

**Movie S4:** Laser ablation of Col-0 emitter facing a Col-0 receiver root with CT-OG FLIM pH probe.

**Movie S5:** H^+^ treatment via OEIP of a R-GECO1 root.

**Movie S6:** Growth of Col-0 root during OEIP treatment.

## Supplemental Information

**Supplemental Information SI1 :**

The Electrochemistry module of COMSOL Multiphysics 6.2**^57^**was used to investigate and visualise the proton delivery. The Tertiary Current Distribution interface includes the Nernst-Planck equation, and can be utilised to find the distribution of different ionic species (*i*) and potential gradients (*V*) as follows:

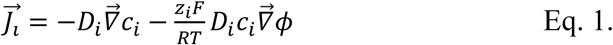

Where *J*_i_ is the flux, *D*_i_ is the diffusion coefficient, *c*_i_ is the concentration, *z*_i_ is the charge of the ionic species, *F* is the Faraday constant, *R* is the gas constant, *T* is the temperature, *ϕ* is the electric potential, and *u* is the bulk flow. An electroneutrality assumption (∑z_i_c_i_ ≈ 0) was used and the convective term was neglected in line with recommendations from Cherian et al. (2023)**^58^**and Handl et al. (2024)**^59^**. The ion composition was reduced to include H^+^, Na^+^, and Cl^-^. As these ions, H^+^ in particular, are known to demonstrate high mobility in highly conductive membranes, no reduced mobility was implemented in any domain. Based on the membrane material mixture, we expect high ion exchange capacity that will enable high fixed charge concentration along with the counteracting high swelling ability. Based on previous analyses of similar materials, we expect the fixed charge concentration to reach approx. 500 mM**^60,61^**. The full parameter list can be found in Table. S**I1**. The initial phase of the experiment, which is characterised by rapid changes **due to large concentration gradients between a fully loaded device and the high pH of the target environment**, is challenging to numerically solve. Therefore, a stepwise method was used. First, steady-state simulations were conducted to find numerical solutions for the initial loaded state. Next, the steady-state solution for the source and membrane in the loaded state was combined with the addition of fresh electrolytes (60 mM NaCl, 1 µM HCl) to shape the initial values for the time-dependent models. To enable convergence of the model during this transition, the device was loaded with NaCl present in the target electrolyte. To further balance H^+^/Na^+^ in the membrane, a current of 400 nA was applied during the virtual device loading. This enabled a solution that involved high H^+^ levels relative to Na^+^ levels (98% vs. 2**%).** The two-minute proton concentration stabilisation phase, which preceded the experiment, can be seen in Figure **SI2**. The final values from this experiment serve as the initial values for the final experiment, **10 min resting followed by 10 min of delivery (Extended data 5a-c**).

**Table SI1:**
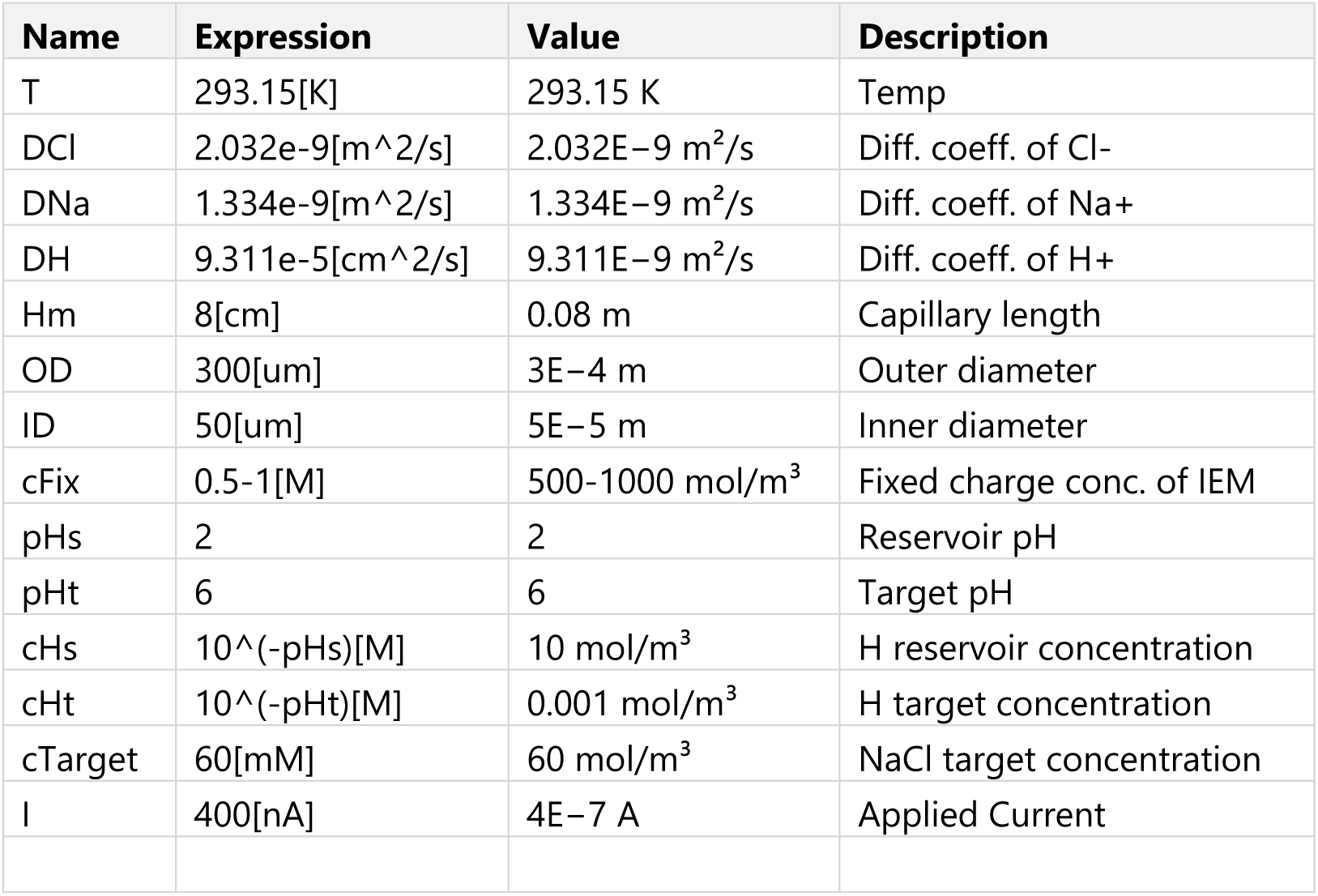
Parameters used in the computational model.

**Figure SI1:**
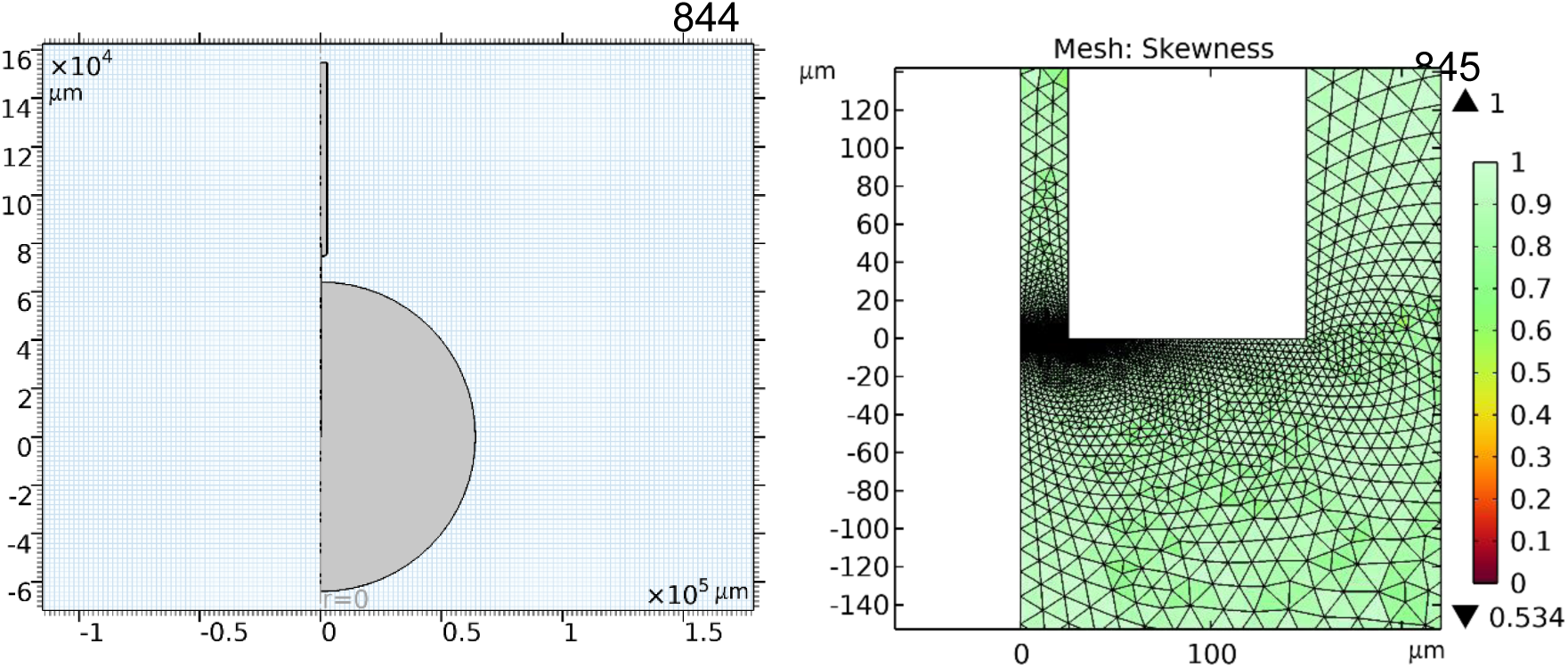
Overview of the geometries used in the computational model. The presented model is a 2D axisymmetric model which is rotated around the z-axis (r=0). The mesh sequence generates a sequence in which the smallest elements are near the membrane interfaces, with the elements gradually increasing further away from these boarders. The model has a total of 21000 elements, with an average quality of 0.85 (skewness).

**Figure SI2:**
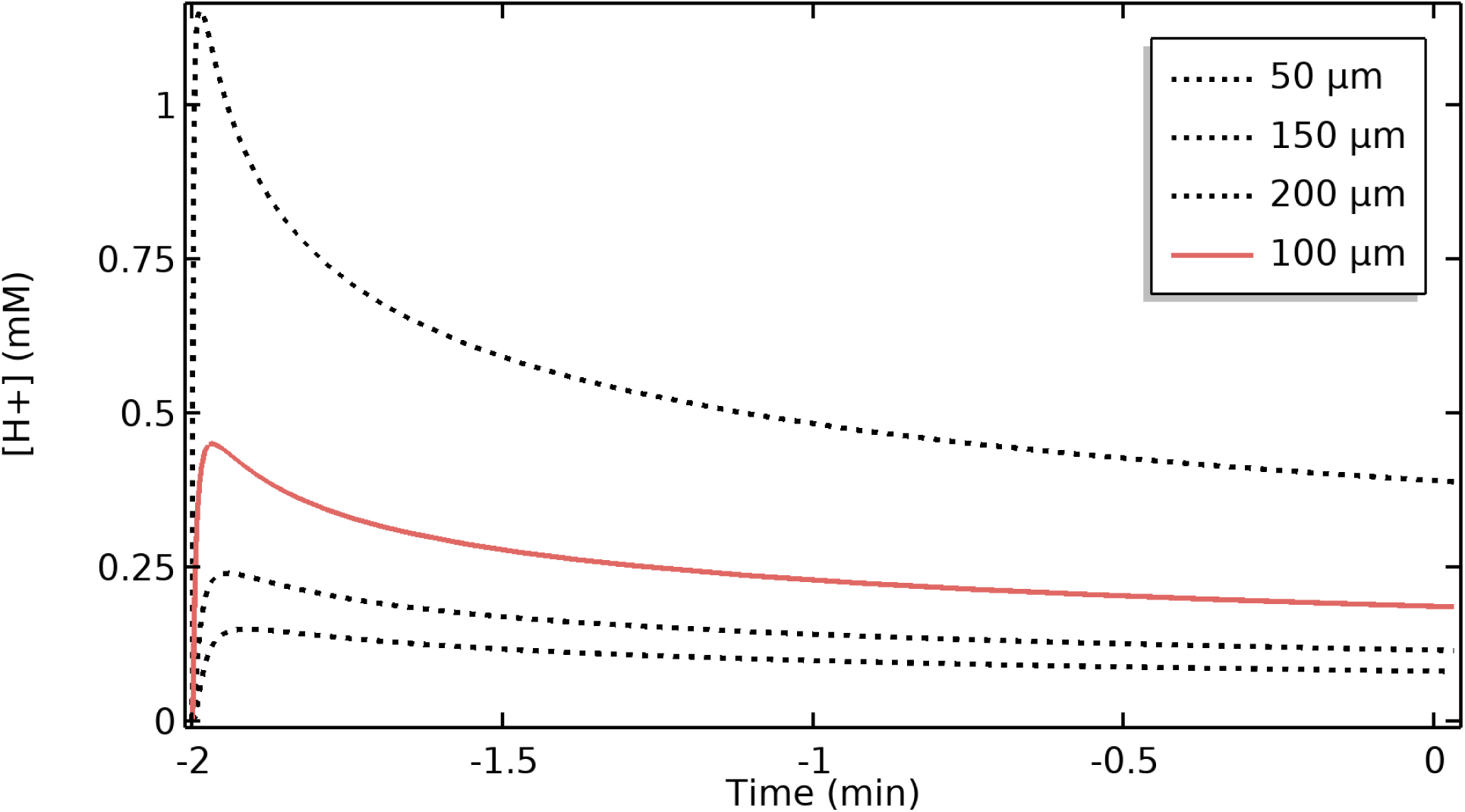
Proton concentrations at distances of 50-200 µm from the device outlet during initial leakage stabilisation. The initial values are from the steady state experiment, the results of which are presented in Fig SI2. The final values from this time series are used as the initial values for the final experiment, with the results discussed in the main text.

**Supplemental information 2 :** The lifetime values obtained with the FLIM-based pH probe CT-OG, were interpolated from the reference values from Besten et al., 2024. The references were plotted, and the third-degree cubic polynomial curve gave the equation parameters required for the interpolation calculation (Fig. SI2). This was done to get the closest representation to the pH values with the most fairly method.

**Fig. SI2:**
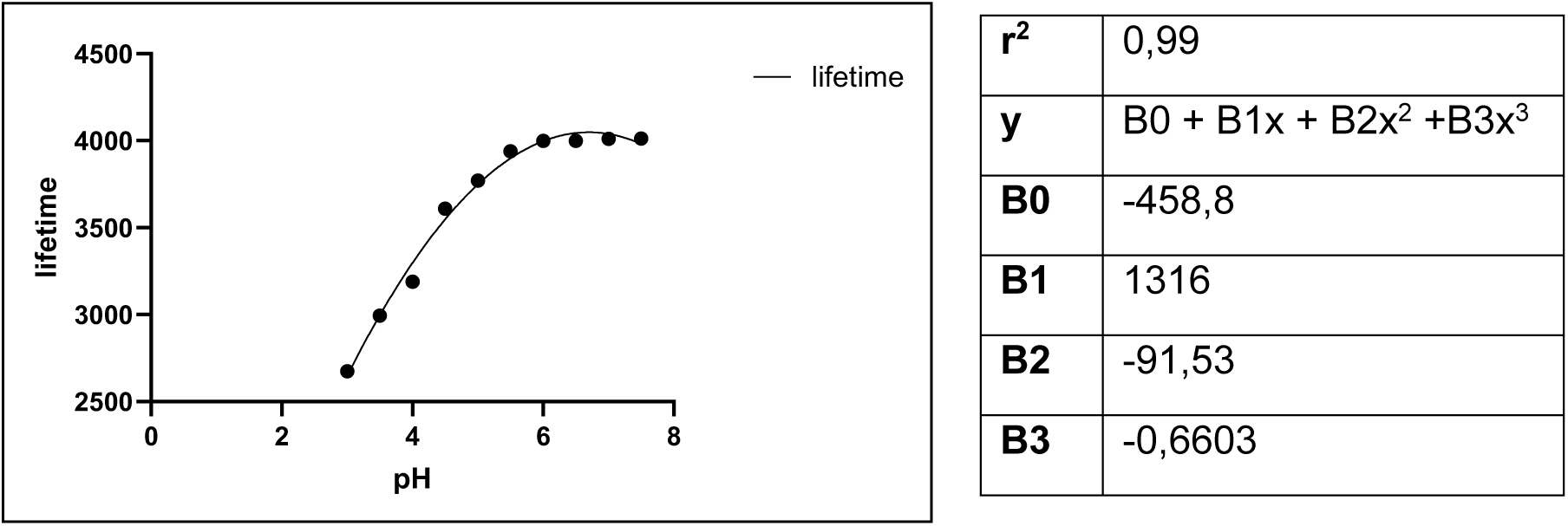
lifetime measured at different pH used as reference for future interpolation calculations. Third degree cubic polynomial regression was applied to obtained equation and parameters essential for future pH values interpolation.

The following script was written in R Studio to calculate the interpolated pH value based on the previous parameters obtained :

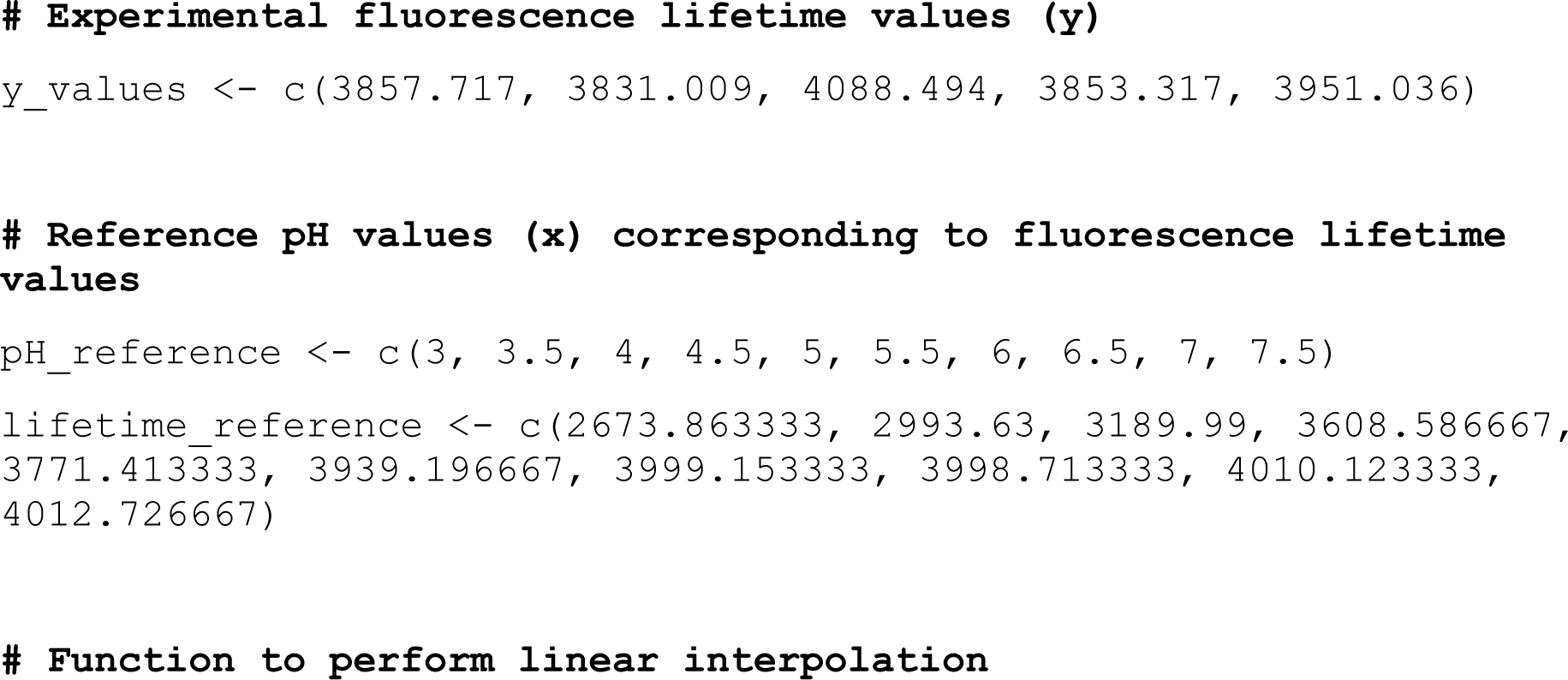

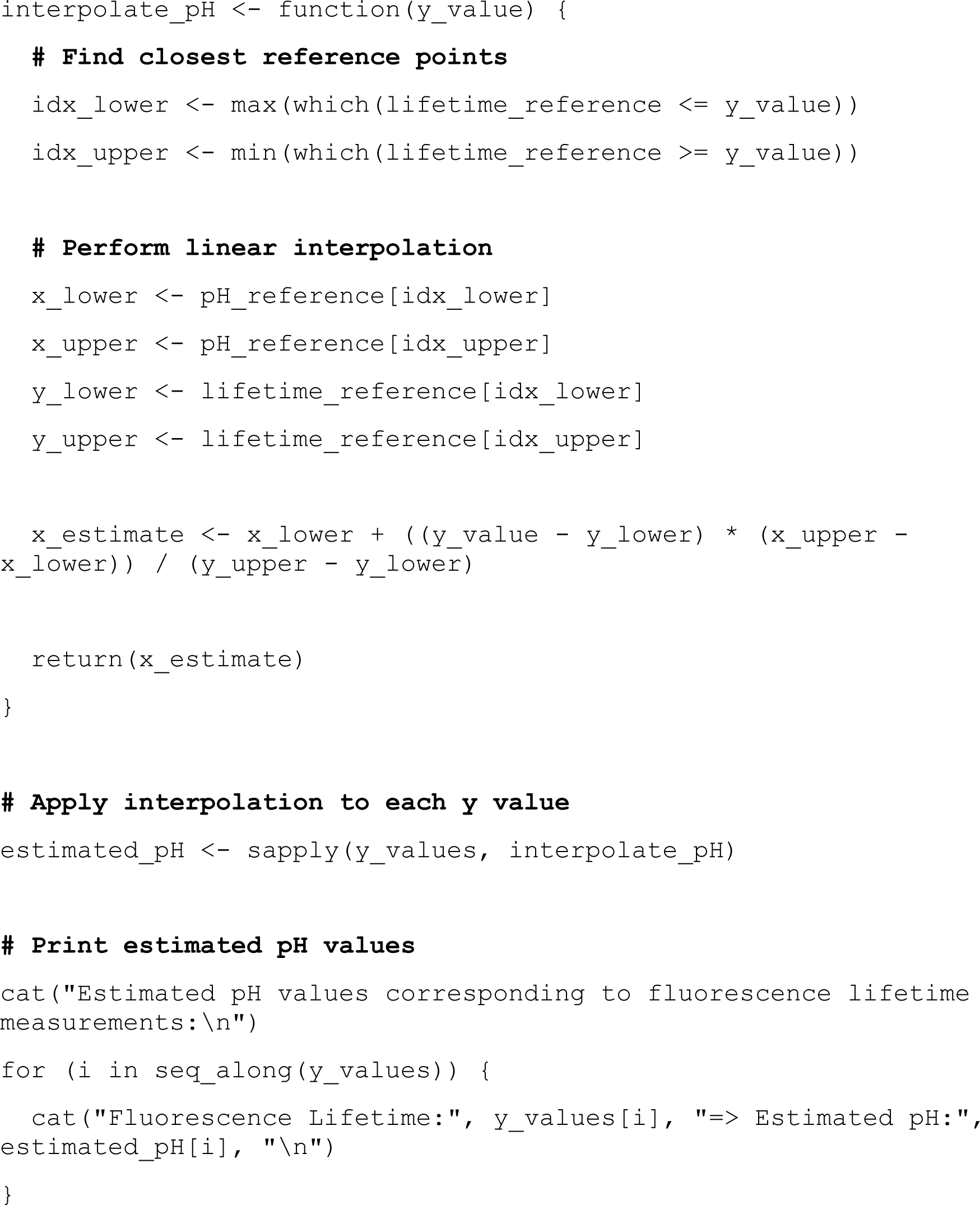

**Supplemental Information 3 :** To obtain the polar histogram that represents the root angle sin the avoidance essay, the following script was written in R Studio :

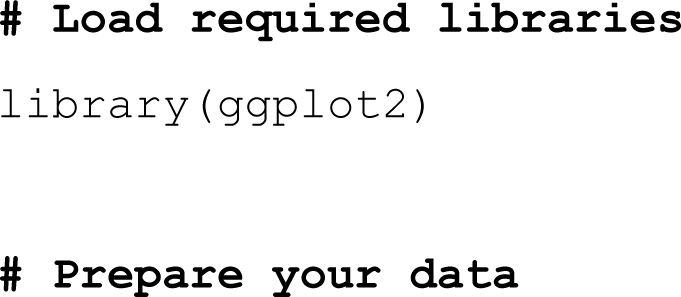

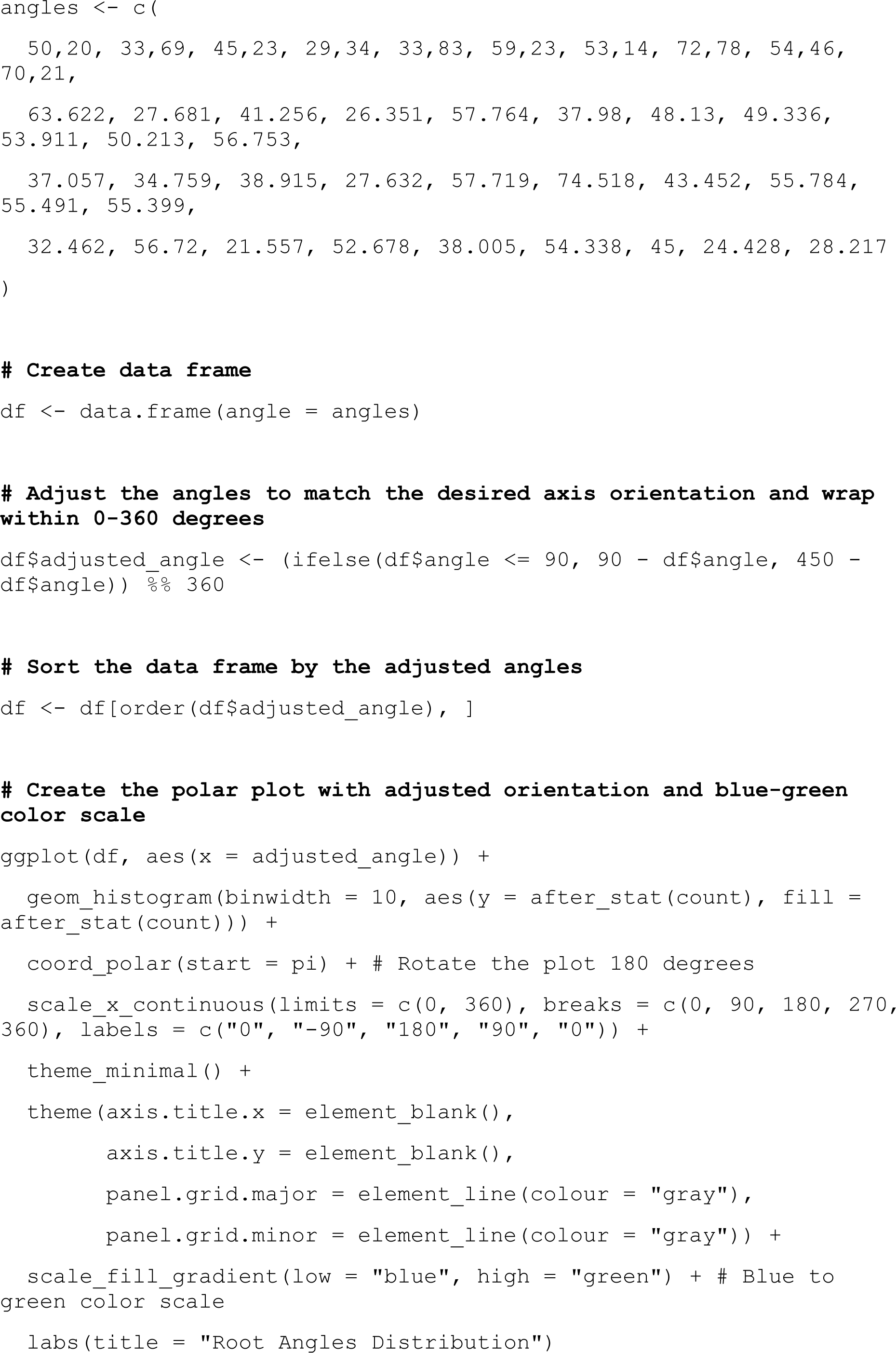

